# Activation of prefrontal parvalbumin interneurons ameliorates treatment-resistant working memory deficit even under continuous antipsychotic treatment in a mouse model of schizophrenia

**DOI:** 10.1101/2023.02.27.530344

**Authors:** Yosefu Arime, Yoshito Saitoh, Mikiko Ishikawa, Chikako Kamiyoshihara, Yasuo Uchida, Kazuki Fujii, Keizo Takao, Kazufumi Akiyama, Noriaki Ohkawa

## Abstract

**BACKGROUND:** One of the critical unmet medical needs in schizophrenia is a remedy for cognitive deficits. However, the neural circuit mechanisms of them remain unresolved. In addition, despite the patients with schizophrenia cannot stop taking antipsychotics due to a high rate of discontinuation-induced relapse, previous studies using animal models of schizophrenia have not considered these clinical situations.

**METHODS:** Here, we employ multi-dimensional approaches, including histological analysis in the prelimbic cortex, LC-MS/MS-based in vivo dopamine D2 receptor occupancy analysis for antipsychotic drugs, in vivo calcium imaging and behavioral analyses of mice using chemogenetic manipulation, to investigate neural mechanisms and potential therapeutic interventions for working memory deficit in a mouse model with chronic phencyclidine (PCP) administration that resembles the schizophrenia symptomatology.

**RESULTS:** Chronic PCP administration led to abnormalities in excitatory and inhibitory synapses, including dendritic spines of pyramidal neurons, vesicular glutamate transporter 1 (VGLUT1) positive terminals, and parvalbumin (PV) positive GABAergic interneurons, in layer 2–3 of the prelimbic cortex. Continuous olanzapine, which achieved a sustained therapeutic window of dopamine D2 receptor occupancy (60–80%) in the striatum, did not affect these synaptic abnormalities and working memory deficit in the PCP-treated mice. We found that the selective prelimbic PV activation, using hM3D(Gq)-DREADD system confirmed by in vivo calcium imaging, restored working memory deficit, even under continuous olanzapine treatment.

**CONCLUSIONS:** Our study raises a possibility that intervention in prefrontal PV neurons leads to an add-on therapy to antipsychotics targeting amelioration of treatment-resistant cognitive deficits in schizophrenia.

## Introduction

One of the critical unmet medical needs in schizophrenia is an effective treatment for cognitive deficits (1). A therapeutic intervention for deficits in cognitive functions, including working memory and executive function, is challenging due to the following reasons: 1) predictive for poor clinical (2) and long-term functional outcomes (3); 2) very high frequency (4); and 3) commonly stable over life-time (5, 6). Unfortunately, cognitive deficits remain largely refractory to existing medications including antipsychotics (7, 8). Hence, novel pharmacotherapies that remedy deficits in patients with schizophrenia are urgently needed. Several meta-analyses have been conducted regarding potential cognitive enhancers targeting various neurotransmitters, including glutamatergic, GABAergic, cholinergic, serotonergic, noradrenergic, and dopaminergic systems (9, 10). These recent literatures showed that the effects of existing cognitive enhancers are little or very small against overall cognition and cognitive subdomains such as working memory and executive function. These clinical data highlight the necessity to understand neural circuit pathology specific to cognitive deficits in schizophrenia and to develop circuit mechanism-based pharmacotherapies for them.

To address the above problem, it is of great advantage to analyze animal models generated by schizophrenia-relevant perturbation. In human, chronic phencyclidine (PCP) use induces a wide range of schizophrenia-like symptomatology including positive and negative symptoms, and cognitive deficits (11-14). In rodents, repeated administration of PCP induces various behavioral phenotypes resembling those of schizophrenia: hypersensitivity to acute PCP treatment, impaired certain types of sociability, and deficits in cognitive functions such as executive function and working memory (15-19). Therefore, chronic PCP-treated rodents have been used as a well-established pharmacological model of schizophrenia (20). We have previously shown that the prelimbic cortex (PL) of the medial prefrontal cortex (mPFC), especially layer 2–3, is a candidate brain region responsible for working memory deficit in chronic PCP-treated mice (18). The structural and functional importance of the PFC in working memory has attracted considerable attention both in humans (21) and rodents (22), and several studies implied impairments of the PL in rodent models of schizophrenia (18, 19). However, the pathological neural circuits that could be definitive targets for improving working memory deficit in this mouse model are still not elucidated.

There are, however, some translational gaps between basic research using animal models and the clinical reality of schizophrenia. First, in real life situations in schizophrenic patients, antipsychotic medication is indispensable for not only improving positive symptoms in the acute phase but also preventing relapse in the maintenance phase (23). All current pharmacological treatments for schizophrenia are dopamine D2 receptor antagonists. PET studies have proposed that an optimal therapeutic window with antipsychotics can be yielded by 60–80% occupancy of striatal dopamine D2 receptors (24, 25). As mentioned above, antipsychotics are effective for positive symptoms but little or none for cognitive deficits. Therefore, cognitive deficits in patients with schizophrenia needs to be ameliorated under the control of psychotic symptoms with antipsychotic treatment. Second, whereas the half-life of antipsychotics in humans is usually 12 to 24 hr, their half-life in rodents tends to be 2 to 4 hr (26). Thus, there is a translational gap between animal models and clinical situations in terms of pharmacokinetic and pharmacodynamic rationale. Despite accumulation of research using animal models of schizophrenia, the neurobiological basis underlying cognitive deficits in schizophrenia remains poorly understood and unaddressed under continuous antipsychotic treatment.

To overcome these problems and bridge the translational gaps, we aimed to investigate a circuit-based therapeutic approach to restore working memory deficit in chronic PCP-treated mice, even under the concurrent continuous antipsychotic treatment. We first examined synaptic pathology, including dendritic spines, excitatory and inhibitory synapses including parvalbumin (PV)-positive interneurons, in the PL associated with working memory deficit in chronic PCP-treated mice. We then conducted manipulation of the PV interneurons using chemogenetic technique, designer receptors exclusively activated by designer drugs (DREADDs), an effect of which was confirmed by in vivo calcium imaging. Also, the neural circuit perturbation in the PL induced prominent effect in amelioration of working memory deficit in chronic PCP-treated mice. In addition, we established antipsychotic dose regimen that could maintain clinically relevant levels of striatal dopamine D2 receptor blockade in mice for several weeks using LC-MS/MS-based in vivo receptor occupancy analysis. Finally, we investigated whether specific PV activation in the PL ameliorate working memory deficit in this mouse model, even under clinically comparable antipsychotic treatment.

## Methods and Materials

### Animals

All animal experiments were performed with the protocol approved by the Ethics Committee of Dokkyo Medical University (Permit Number: 0704, 0771, and 1112, and University of Toyama (A2019OPR-1, and A2019OPR-2), performed in accordance with the Guidelines for Care and Use of Laboratory Animals, and conformed to all Japanese federal animal welfare rules and guidelines. Thy1-GFP line O (Stock No. 007919) and PV-Cre (Stock No. 017320) mice were purchased from the Jackson Laboratory. C57BL/6J mice were obtained from Japan Clea Co (Key Resources Table). Strains were backcrossed to congenic C57BL/6J mice. All experiments used male mice. See Key Resources Table for strain information.

Detailed methods and materials regarding additional animals, drug, viral vector preparation, surgery and drug treatment, histological experiments, behavioral tests, in vivo calcium imaging, and LC-MS/MS experiments are described in the Supplemental Methods.

## Results

### Chronic PCP administration led to synaptic abnormalities in the PL

We have previously shown that PL, especially layer 2–3, is a candidate brain region responsible for working memory deficit in chronic PCP-treated mice (18). However, the circuit mechanism that can be a therapeutic target is unclear. We first examined pathological basis for working memory deficit in this region of chronic PCP-treated mice. Previous studies show that the size of dendritic spine head is tightly coupled to synaptic function such as the size of postsynaptic density, the number of AMPA receptors, and the synaptic strength (27-29). To determine the effect of chronic PCP treatment on spine pathology, we reconstructed three-dimensional dendritic segments using Thy1-GFP line O mice (30) and then performed high-throughput, detailed, and cell-specific dendritic spine analysis with NeuronStudio (31). The laminar cytoarchitecture in the PL was identified by Ctip2 (a layer 5 marker) and Foxp2 (a layer 6 marker) immunostaining (Figure 1B). We found that chronic PCP administration led to decreases in total (Figure 1C, left) and each type of spine density (Figure 1C, middle) on pyramidal neurons in layer 2–3 of the PL. Analysis of frequency distribution of head diameter of dendritic spines (Figure 1C, right), head diameter (Supplemental figure S1A) and spine length of each spine type (Supplemental figure S1B) revealed no changes in spine proportions, suggesting that chronic PCP reduced dendritic spine density regardless of spine types in this region. Both in layer 5 and 6, no differences were observed in spine density (layer 5: Figure 1D left and middle, layer 6: Figure 1E left and middle), head diameter (layer 5: Figure 1D right, layer 6: Figure 1E right, Supplemental Figure S1C–F) between chronic saline-and PCP-treated mice. These data from detailed-and layer-specific dendritic spine analysis corroborate and extend data from previous study using electron microscopy (32). Together, these results suggest that chronic PCP-treated mice have abnormalities in receiving of excitatory synaptic inputs to layer 2–3 pyramidal neurons of the PL.

**Figure 1.**
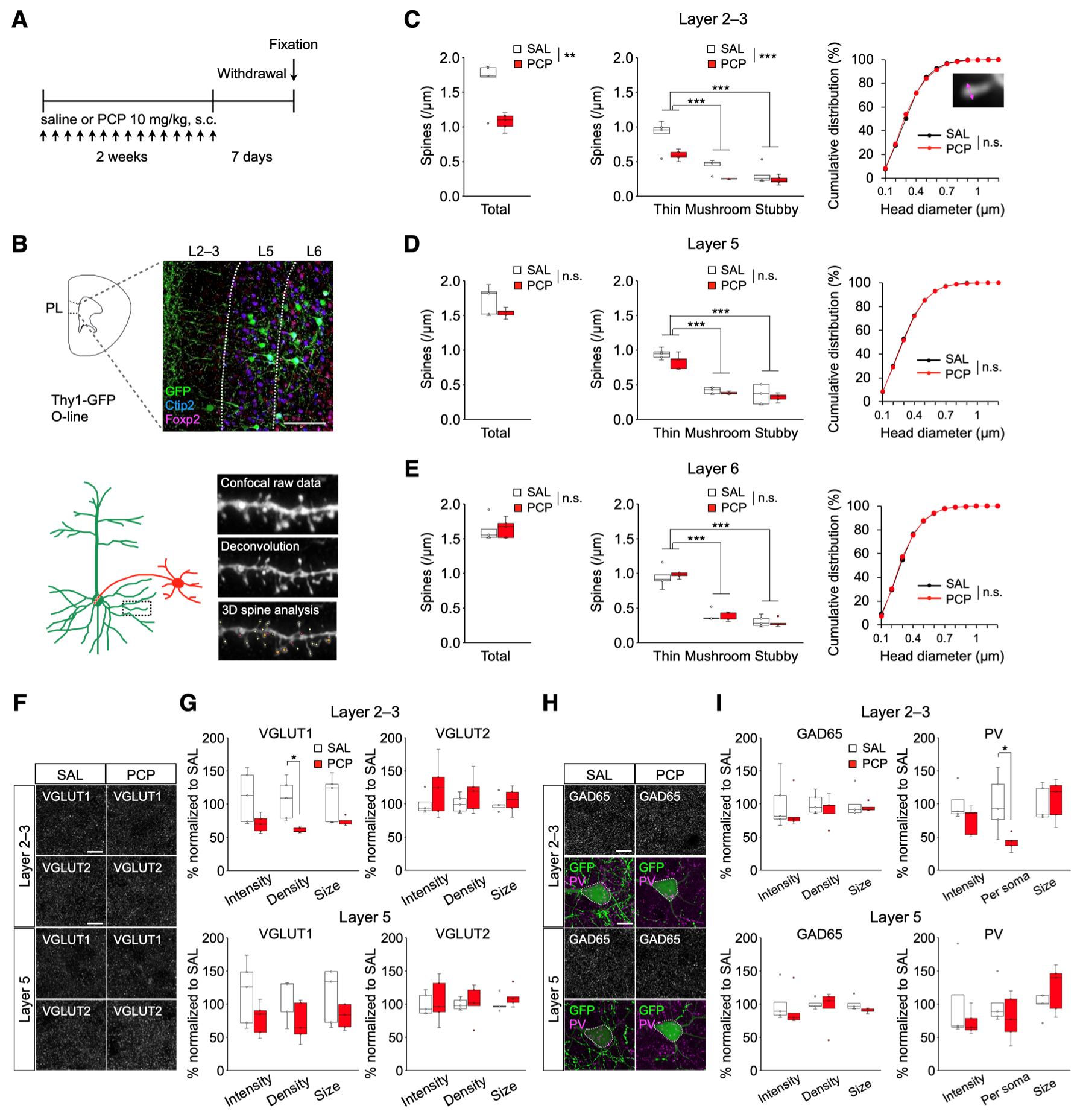
Selective loss of dendritic spines and abnormalities of excitatory and inhibitory synapses in layer 2–3 of the PL of chronic PCP-treated mice. **(A)** Schematic timeline for morphological analysis. **(B)** Representative confocal images in the PL before and after 3D deconvolution, and 3D reconstructions of dendritic spines by NeuronStudio. Scale bar: 100 µm. **(C–E)** Dendritic spine analysis in the PL. In layer 2–3 of the PL **(C)**, chronic PCP administration results in decreases in total (left) and each spine density (middle). In layer 5 **(D)** and 6 **(E)**, no changes were observed in total (left) and each spine density (middle). Cumulative frequency distributions of overall spine head diameter in layer 2–3 **(C, right)**, 5 **(D, right)**, and 6 **(E, right)** are plotted. There are no changes in the distribution of spine head size in all layers. **(F)** Representative images of immunostaining for VGLUT1 and VGLUT2 in the PL of chronic saline-or PCP-treated mice. Scale bars: 10 µm. **(G)** Quantification of VGLUT1 and VGLUT2 positive puncta. Decreased density of VGLUT1 positive puncta in layer 2–3 of the PL in chronic PCP-treated Thy1-GFP mice were observed (top, left). **(H)** Representative images of immunostaining for GAD65 and PV in the PL of chronic saline-or PCP-treated Thy1-GFP mice. Scale bars: 10 µm. **(I)** Quantification of GAD65 and PV positive puncta. Decreased PV positive puncta surrounding pyramidal neuron soma in layer 2–3 of the PL in chronic PCP-treated Thy1-GFP mice were observed (top, right). In box plots, the central mark indicates the median and the bottom and top edges of the box indicate the 25th and 75th percentiles, respectively. n.s., no significance. \*\**p* < 0.01, \*\*\**p* < 0.001.

Next, we tested whether chronic PCP could affect excitatory and inhibitory pre-synapses in the PL. Neurons in the cerebral cortex receive two main types of excitatory inputs, that is, vesicular glutamate transporter 1 (VGLUT1) is predominantly localized within the terminals of cortical projections and VGLUT2 is localized within the terminals of thalamus and hypothalamus (33). Excitatory input sources were identified by immunostaining with anti-VGLUT1 or VGLUT2 antibodies to distinguish cortico-cortical or thalamo-cortical inputs, respectively (34). We found that chronic PCP administration led to decrease in VGLUT1+ puncta density in layer 2–3 of the PL with no change in layer 5 (Figure 1G, left). Both in layer 2–3 and layer 5, no differences were observed in VGLUT2+ puncta between chronic saline-and PCP-treated mice. (Figure 1G, right). Together, our results demonstrate that cortical excitatory inputs to layer 2–3 of the PL were specifically attenuated in this PCP model. Previous studies reported chronic ketamine, a PCP analog, reduced PV neuron density in the mPFC (19). To examine whether inhibitory synaptic inputs in the PL are affected by chronic PCP administration, we assessed inhibitory synapses by quantifying puncta of glutamic acid decarboxylase 65 (GAD65), as a marker of GABAergic neuron terminals, and PV-immunoreactive puncta surrounding pyramidal neuron soma, presumed PV+ basket cell terminals (35). We found that chronic PCP administration led to decrease in PV+ puncta surrounding GFP+ pyramidal neuron cell bodies in layer 2–3 of the PL (Figure 1I, right). Both in layer 2–3 and layer 5, no differences were observed in GAD65+ puncta between chronic saline-and PCP-treated mice. (Figure 1I, left). Overall, these anatomical data revealed that PV+ puncta density onto pyramidal neuron soma was reduced in the layer 2–3 of the PL, suggesting an attenuated inhibitory inputs from PV+ basket cells in chronic PCP-treated mice.

### Chemogenetic activation of prefrontal PV neurons restores working memory deficit in chronic PCP-treated mice

Because our histological data indicate chronic PCP-treated mice exhibited deficits in PV neuron-mediated excitatory and inhibitory circuits in the PL, we next examined whether prefrontal PV neuron activation could restore working memory deficit in this mouse model. We first applied hM3D(Gq), the Gq-coupled excitatory DREADD, and G-CaMP7 to PV interneurons by AAV-hSyn-DIO-hM3D(Gq)-mCherry and AAV-hSyn-DIO-G-CaMP7 coinjection into the PL of PV-Cre mice (Figure 2A, left). After implantation of a GRIN lens in the PL, we imaged the calcium signals of PV neuron during pretreatment, vehicle and deschloroclozapine (DCZ) treatment with a Nipkow-disk confocal microscopy (Figure 2A, right and 2B). Chemogenetic perturbation has been used as a powerful tool for manipulating of neuronal activity and behaviors. In using this technique, we noticed recent studies that clozapine-N-oxide (CNO), widely used as a DREADD agonist, does not enter the brain after systemic administration, and rapidly converted to clozapine, clozapine readily enters the brain, and activates DREADDs, leading to potential off-target effects (36). On the other hand, a novel DREADD actuator DCZ is an extremely potent, highly brain penetrant, metabolically stable and selective actuator for hM3D(Gq) and hM4D(Gi) DREADDs in mice (37). To minimize off-target effects and to increase specificity of PV neurons, we employed a novel approach for increasing prelimbic PV neuron activity using an excitatory DREADD and its ligand DCZ.

**Figure 2.**
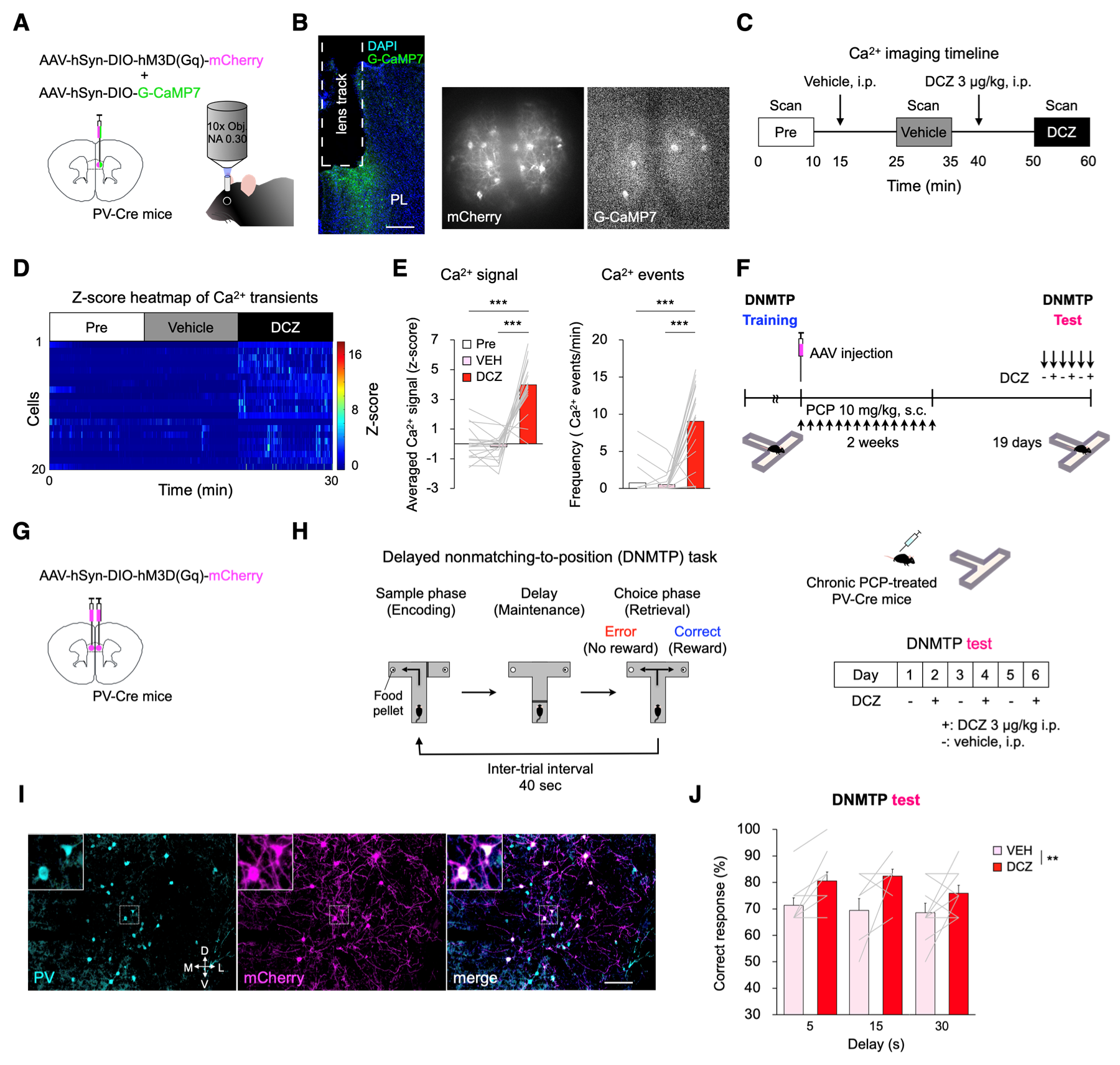
Chemogenetic activation of PV neurons in the PL improves working memory deficit in chronic PCP-treated mice. **(A)** Schema of viral delivery of AAV-hSyn-DIO-G-CaMP7 and AAV-hSyn-DIO-hM3D(Gq)-mCherry to PL (left). Schematic of imaging of PV neuron activity in the PL of an awake mouse (right). **(B)** GRIN lens implantation (dashed line) into the PL. Scale bar: 300 µm (left). Representative mCherry (middle) and stacked G-CaMP7 (right) images acquired through Nipkow-disk confocal microscopy. **(C)** Schematic timeline of confocal scanning. (**D**) Z-score heatmap of average Ca^2+^ transients during pretreatment, vehicle and DCZ 3 µg/kg treatment. **(E)** Comparison of z-score of average Ca^2+^ signals (left) and Ca^2+^ event frequency (right) in 10 min bins. DCZ 3 µg/kg treatment significantly increased both averaged Ca^2+^ signals and frequency of Ca^2+^ event of PV neuron. Data are presented as the mean. Each line represents each cell. \*\*\**p* < 0.001. **(F)** Schematic timeline of DNMTP test with T-maze. **(G)** Schema of bilateral viral delivery of AAV-hSyn-DIO-hM3D(Gq)-mCherry to the PL. **(H)** Schematic illustration of test sessions in DNMTP task with T-maze (left). Test sessions of DNMTP task consist of six days with DCZ 3 µg/kg every other day (right). **(I)** Representative images of PV immunostaining (cyan) and viral expression (magenta) in the PL. Scale bar: 100 µm. **(J)** DCZ 3 µg/kg significantly increased the percent correct responses across all delay periods in chronic PCP-treated PV-Cre mice that previously received AAV-hSyn-DIO-hM3D(Gq)-mCherry (right). Data are presented as the mean ± SEM. Each line represents each mouse. \*\**p* < 0.01.

We confirmed increased Ca^2+^ signal intensities and frequency of Ca^2+^ events of PV neurons after systemic DCZ 3 µg/kg treatment, suggesting prelimbic PV neuron activities were enhanced by hM3D(Gq)-DREADD (Figure 2C–E, Supplemental figure S2). To determine whether the prelimbic PV neuron activation restore working memory deficit, PV-Cre mice that were injected with AAV-hSyn-DIO-hM3D(Gq)-mCherry into the bilateral PL were tested after the completion of chronic PCP treatment (Figure 2F, G). We virally targeted hM3D(Gq)-mCherry to PV neurons in the PL of PV-Cre mice prior to chronic PCP administration (Figure 2I). Compared with vehicle injection, DCZ 3 µg/kg led to a significant increase in percentage of correct responses across all delay periods in chronic PCP-treated PV-Cre mice that previously received AAV-hSyn-DIO-hM3D(Gq)-mCherry (Figure 2J). The response latencies to reach pellet cup did not differ regardless of DCZ treatment (Supplemental figure S3A) or behavioral responses (Supplemental figure S3B), suggesting that improved performance reflects improved working memory. These results demonstrate that chemogenetic activation of prelimbic PV neurons improve working memory deficit in chronic PCP-treated mice.

### LC-MS/MS-based in vivo dopamine D2 receptor occupancy with continuous antipsychotic treatment

We determined whether continuous haloperidol (typical antipsychotic drug) and olanzapine (atypical antipsychotic drug) treatment could reach and maintain dopamine D2 receptor occupancy (60–80%), which is an indicator for the clinical situation, for several weeks in mice. The aim of this experiments is twofold: (1) to establish antipsychotic doses that could maintain clinically relevant levels of striatal D2 receptor blockade in mice for several weeks, and (2) to apply as a tool to validate their ability to inhibit behavioral abnormalities, including working memory deficit, in chronic PCP-treated mice. We performed in vivo receptor occupancy assays in the mouse striatum for dopamine D2 receptors (Figure 3A, details see in Supplemental Methods and Supplemental Table S2–4). We continuously administered haloperidol or olanzapine to mice for a maximum of 4 weeks (Figure 3B). In vivo D2 receptor occupancy analysis revealed that haloperidol 0.5 mg/kg/day achieved and maintained clinical levels of dopamine D2 receptor occupancy (∼60–80%) for at least 13 days (Figure 3C) and olanzapine 7.5 mg/kg/day did those levels for at least 28 days (Figure 3D, Supplemental Table S5).

**Figure 3.**
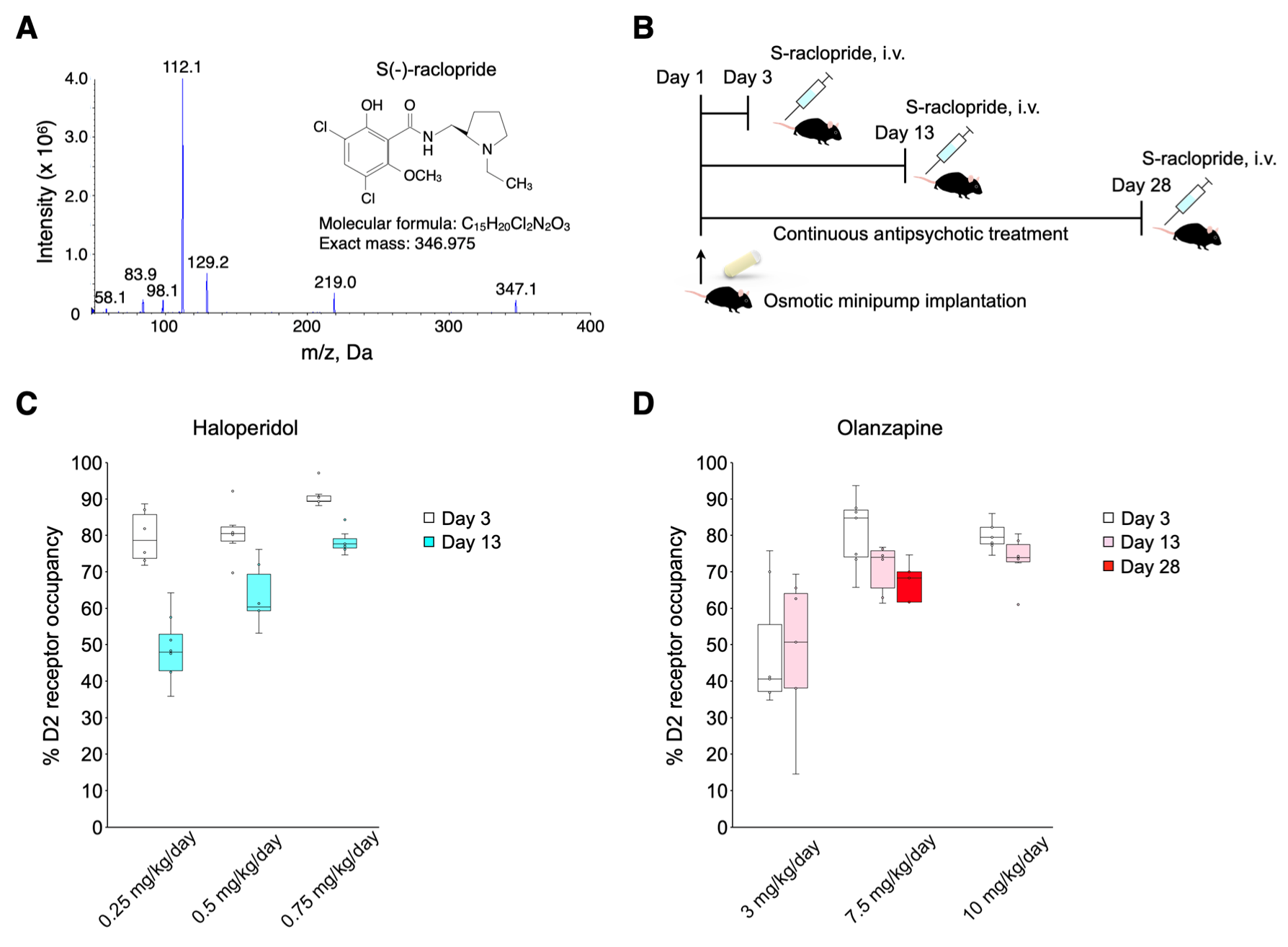
In vivo striatal dopamine D2 receptor occupancy during continuous antipsychotic treatment in mice. **(A)** Fragmentation pattern of S(-)-raclopride during product ion scan. **(B)** Schematic timeline of continuous antipsychotic treatment with osmotic minipump. **(C)** D2 receptor occupancy levels produced by haloperidol, a typical antipsychotic drug. **(D)** D2 receptor occupancy levels produced by olanzapine, an atypical antipsychotic drug. Clinical levels of dopamine D2 receptor occupancy (60–80 %) were achieved with haloperidol 0.5 mg/kg/day for 2 weeks and olanzapine 7.5 mg/kg/day for 4 weeks. In box plots, the central mark indicates the median and the bottom and top edges of the box indicate the 25th and 75th percentiles, respectively.

### Effects of clinically relevant antipsychotic dose regimen on histological and behavioral abnormalities in chronic PCP-treated mice

Current antipsychotic drugs are relatively effective for positive symptoms but little or none for cognitive deficits (1, 23). Based on these clinical situations, we hypothesized that continuous antipsychotic treatment, which maintain clinical levels of dopamine D2 receptor occupancy in mouse striatum, could ameliorate acute PCP-induced psychomotor behavior, but may fail to improve working memory deficit in chronic PCP-treated mice.

To test this, we first examined whether continuous haloperidol or olanzapine treatments, which maintain clinical levels of dopamine D2 receptor occupancy for at least 2 weeks, would affect acute PCP challenge-induced hyperlocomotion (Figure 4A). The open field test showed repeated administration of PCP developed behavioral sensitization across all groups as well (Supplemental figure S4A-F). After withdrawal from chronic PCP treatment, acute PCP-induced locomotor activity and c-Fos expression in the nucleus accumbens (NAc) was measured. Acute PCP 3 mg/kg challenge induced hyperlocomotion in mice chronically treated with PCP (Figure 4B, C) and increased the number of entries and distance traveled both in the corner and center zone (Supplemental figure S5A-D). Only olanzapine 7.5 mg/kg/day attenuated acute PCP-induced hyperactivity (Figure 4C) and reduced the number of entries and distance traveled both in the corner and center zone (Supplemental figure S5A-D). Acute PCP challenge with open field test also increased c-Fos expression in the shell of the NAc (Figure 4D, E). Olanzapine 7.5 mg/kg/day reduced acute PCP-induced c-Fos elevation in this region.

**Figure 4.**
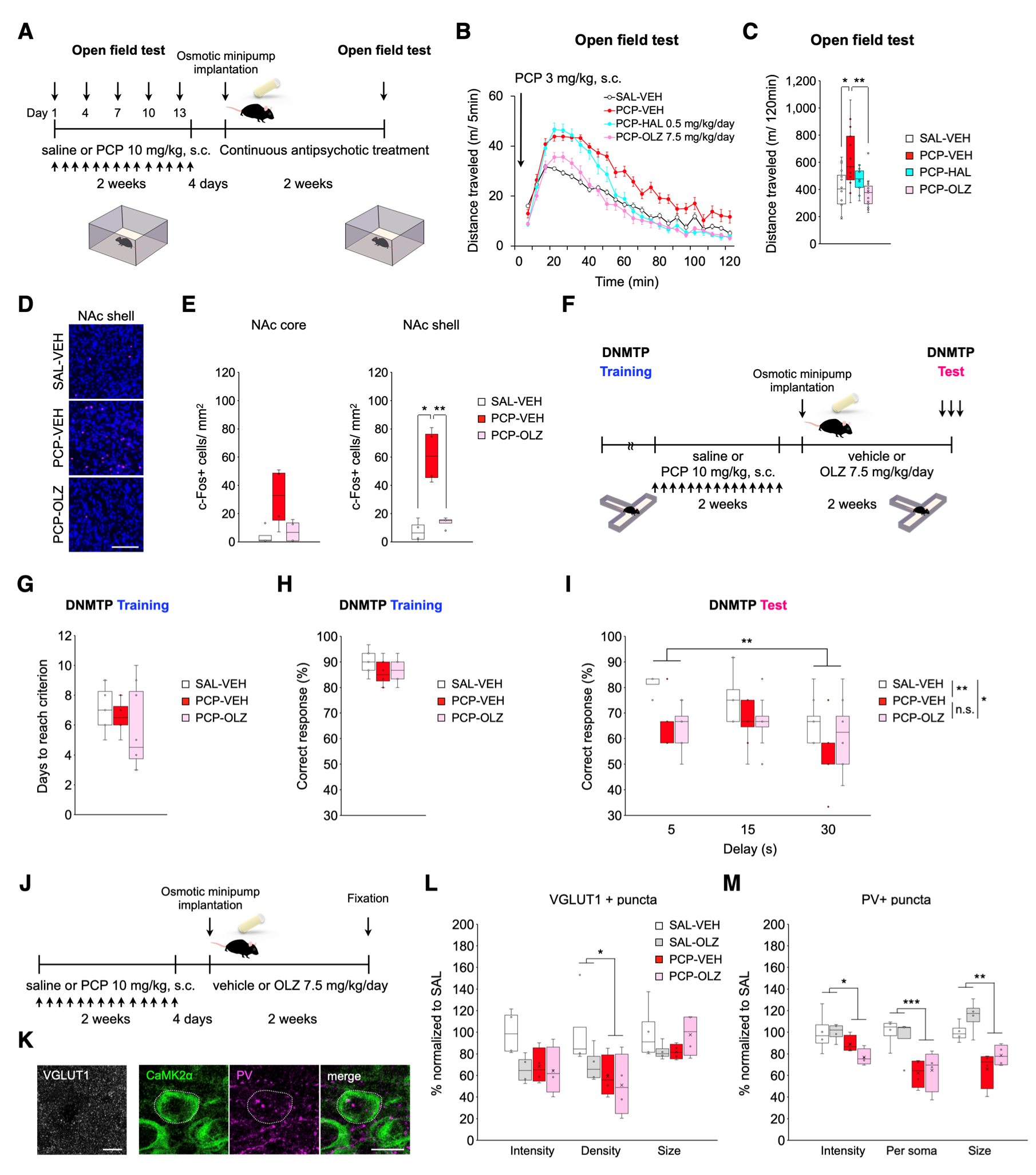
Effects of continuous olanzapine treatment on synaptic and behavioral abnormalities in chronic PCP-treated mice. **(A)** Schematic timeline of open field test procedure. **(B)** Distance traveled in 5 min bins following acute PCP 3 mg/kg challenge in the open field during continuous antipsychotic treatment. **(C)** Acute PCP treatment significantly increased locomotor activity in chronic PCP-treated mice. Compared with continuous vehicle treatment, olanzapine (OLZ) 7.5 mg/kg/day significantly reduced acute PCP-induced locomotor hyperactivity in chronic PCP-treated mice. **(D)** Representative images of c-Fos immunostaining in the shell of the nucleus accumbens (NAc). Scale bar: 100 µm. **(E)** Quantification of c-Fos positive cells in the NAc core (left) and shell (right). Acute PCP treatment significantly increased c-Fos positive cells in the NAc shell of chronic PCP-treated mice. Compared with continuous vehicle treatment, OLZ 7.5 mg/kg/day significantly reduced acute PCP-induced c-Fos elevation in chronic PCP-treated mice. **(F)** Schematic timeline of DNMTP task with T-maze during continuous olanzapine treatment. During training sessions of DNMTP task, no group differences in days to reach criterion **(G)** and correct responses **(H)** were observed before repeated administration of saline or PCP. **(I)** Similar to our previous study, repeated administration of PCP reduced percent correct responses during DNMTP task with T-maze. Consistent with continuous vehicle treatment (PCP-VEH), OLZ 7.5 mg/kg/day (PCP-OLZ 7.5 mg/kg/day) did not affect working memory deficit in chronic PCP-treated mice. **(J)** Schematic timeline of morphological analysis after continuous vehicle or OLZ treatment both in chronic saline-or PCP-treated mice. **(K)** Representative images of immunostaining for VGLUT1 (left), CaMK2a (right, green), and PV (right, magenta). Scale bar: 10 µm. **(L)** Quantification of VGLUT1 positive puncta. Decreased density of VGLUT1 positive puncta in layer 2–3 of the PL in chronic PCP-treated mice were observed. OLZ 7.5 mg/kg/day for two weeks did not affect VGLUT1 positive puncta in layer 2–3 of the PL both in chronic saline-and PCP-treated mice. **(M)** Quantification of PV positive puncta surrounding CaMK2a positive pyramidal neuron soma. Decreased expression of PV (intensity, per soma, and size) in layer 2–3 of the PL in chronic PCP-treated mice were observed. OLZ 7.5 mg/kg/day for two weeks did not affect PV positive puncta in layer 2–3 of the PL both in chronic saline-and PCP-treated mice. In box plots, the central mark indicates the median and the bottom and top edges of the box indicate the 25th and 75th percentiles, respectively. n.s., no significance. \**p* < 0.05, \*\**p* < 0.01, \*\*\**p* < 0.001.

Next, we examined whether continuous olanzapine treatment would affect working memory deficit in chronic PCP-treated mice (Figure 4F). Working memory was assessed using delayed nonmatching-to-position (DNMTP) task with T-maze which has been described previously (18). In DNMTP training, there was no significant group differences in days to reach criterion (Figure 4G) or in correct response rates (Figure 4H) before continuous vehicle or olanzapine treatment. Similar to our previous study (18), repeated administration of PCP reduced percent correct responses during DNMTP task with T-maze (Figure 4I). We found olanzapine 7.5 mg/kg/day had no effect on reduced correct responses in chronic PCP-treated mice. These results showed that clinically comparable olanzapine treatment ameliorate psychomotor behavior and neural hyperactivation in the shell of the NAc but do not affect working memory deficit in our PCP model, suggesting that they share similar poor clinical efficacy of antipsychotics.

We then tested whether continuous olanzapine treatment could affect synaptic abnormalities in layer 2–3 of the PL of chronic PCP-treated mice (Figure 4J). We found that repeated administration with PCP reduced VGLUT1+ (Figure 4L) and PV+ puncta (Figure 4M) and that continuous treatment with olanzapine 7.5 mg/kg/day had no effect on these synaptic abnormalities both in chronic saline-and PCP-treated mice. Previous reports have demonstrated that prefrontal PV+ neuron density was reduced in rats chronically treated with ketamine (19) and patients with schizophrenia (38). Similar to these reports, we observed that repeated administration of PCP reduced PV+ neuron density in the PL approximately three weeks after withdrawal from PCP (Supplemental figure S6A, B). We also found olanzapine 7.5 mg/kg/day had no effect on reduced PV+ neuron density. Together, these results suggest that there are long-lasting abnormalities in prelimbic PV neurons of mice chronically treated with PCP and that continuous olanzapine treatment do not affect excitatory and inhibitory synaptic abnormalities in the PL, especially layer 2–3, as well as working memory deficit in our PCP model.

### Chemogenetic activation of prefrontal PV neurons in combination with clinically comparable olanzapine treatment restores working memory deficit in chronic PCP-treated mice

We examined whether ameliorating effect of working memory by prefrontal PV activation using hM3D(Gq)-DREADD is exerted even under treatment with clinically relevant antipsychotic dose regimen, which reflect medication status in patients with schizophrenia. We applied hM3D(Gq)-DREADD to PV interneurons by AAV-hSyn-DIO-hM3D(Gq)-mCherry injection into the PL (Figure 5A, B). AAV-hSyn-DIO-mCherry was used to control potential effects of DCZ. We virally targeted hM3D(Gq)-mCherry or mCherry to PV neurons in the PL of chronic PCP-treated PV-Cre mice with subsequent olanzapine 7.5 mg/kg/day treatment (Figure 5C). In DNMTP training, there were no significant differences in days to reach criterion (Supplemental figure S7A) or in correct response rates (Supplemental figure S7B) between two types of AAV groups noted above. In DNMTP test sessions under olanzapine 7.5 mg/kg/day treatment, we found DCZ 3 µg/kg led to a significant increase in percentage of correct responses across all delay periods in chronic PCP-treated PV-Cre mice that previously received AAV-hSyn-DIO-hM3D(Gq)-mCherry (Figure 5E, right). Such recovery effect of DCZ was not observed in chronic PCP-treated PV-Cre mice that previously received AAV-hSyn-DIO-mCherry (Figure 5E, left). Response latency did not change with treatment conditions (Supplemental figure S8A) or behavioral responses (Supplemental figure S8B), suggesting that improved performance reflects improved working memory. To further examine the roles of prefrontal PV activation on behavior in PCP model, we performed open field test in chronic PCP-treated PV-Cre mice that previously received AAV-hSyn-DIO-mCherry or AAV-hSyn-DIO-hM3D(Gq)-mCherry under olanzapine 7.5 mg/kg/day treatment. We found that DCZ 3 µg/kg did not change in acute PCP-induced locomotor hyperactivity between hM3D(Gq)-mCherry-and mCherry-expressing mice, suggesting that prelimbic PV activation does not affect inhibitory effects of continuous olanzapine treatment on psychomotor behavior (Supplemental figure S9D, E). Taken together, these results demonstrate that chemogenetic activation of prelimbic PV neurons ameliorate working memory deficit in chronic PCP-treated mice, even under continuous olanzapine treatment without affecting improvement effects of olanzapine.

**Figure 5.**
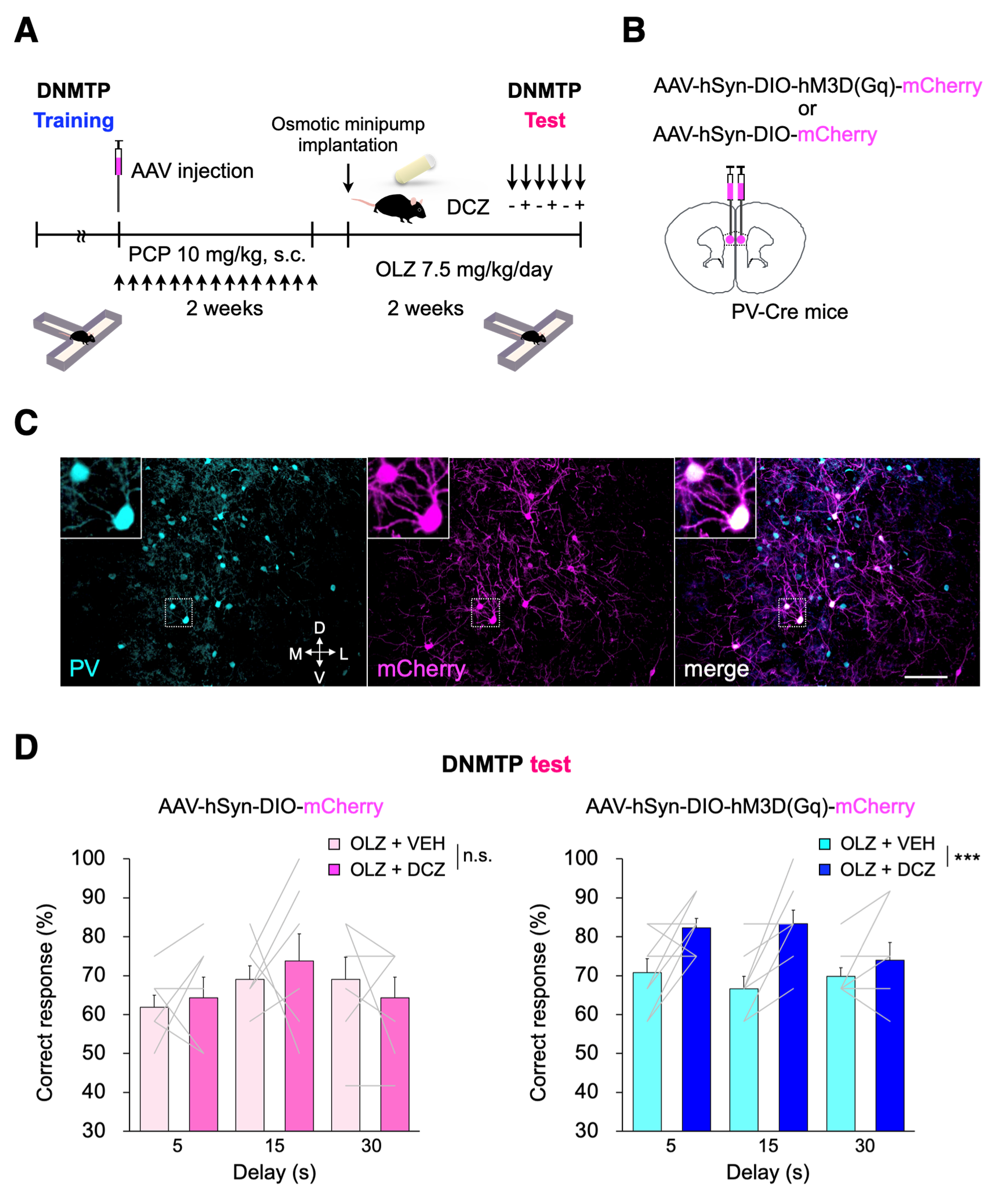
Chemogenetic activation of PV neurons in the PL improves working memory deficit in chronic PCP-treated mice even under continuous OLZ treatment. **(A)** Schematic timeline of DNMTP task with T-maze. **(B)** Schema of bilateral viral delivery of AAV-hSyn-DIO-mCherry or AAV-hSyn-DIO-hM3D(Gq)-mCherry to the PL. **(C)** Representative images of PV immunostaining (cyan) and viral expression (magenta) in the PL. Scale bar: 100 µm. **(D)** The percent correct responses were not changed by DCZ in chronic PCP-treated PV-Cre mice that previously received AAV-hSyn-DIO-mCherry during continuous OLZ treatment (left). DCZ 3 µg/kg significantly increased the percent correct responses across all delay periods in chronic PCP-treated PV-Cre mice that previously received AAV-hSyn-DIO-hM3D(Gq)-mCherry during continuous OLZ treatment (right). Data are presented as the mean ± SEM. Each line represents each mouse. n.s., no significance. \*\**p* < 0.01.

## Discussion

The main findings of our study are that chronic PCP administration leads to working memory deficit, reduced excitatory synapses from other cortex, and attenuated PV neuron terminals onto the pyramidal neuron soma in layer 2–3 of the PL, none of which were rescued by antipsychotic treatment, but that chemogenetic activation of PV neurons in this region ameliorates working memory deficit in PCP mouse model, even under continuous olanzapine treatment. On the other hand, prefrontal PV activation does not affect the therapeutic effects of continuous olanzapine on acute PCP-induced hyperactivity. Our findings using pharmacological model of schizophrenia support the hypothesis that abnormal balance between excitatory and inhibitory tones in the PFC is implicated in the pathophysiology of cognitive deficits in schizophrenia (39, 40) and the findings herein suggest that prefrontal PV neuron activation may prove to be a promising remedy for working memory deficit in schizophrenia.

Numerous cognitive tasks, including the novel object recognition task and other forms of cognitive tasks, have been widely used in preclinical studies. Although existing antipsychotics have minimal or even adverse effects on cognitive deficits in the clinical situation, many preclinical studies have shown that current antipsychotics restore deficits in these tasks, suggesting they have inconsistent results with clinical effects and will have limited predictive validity (41). We found continuous olanzapine treatment, which maintain clinical levels of dopamine D2 receptor occupancy in the mouse striatum for at least 4 weeks, could ameliorate acute PCP-induced psychomotor behavior, but not improve working memory deficit in chronic PCP-treated mice. These results indicate that our PCP model definitely reflects antipsychotics-resistant cognitive deficits. Together, antipsychotics-resistant working memory deficit in our PCP model may be a good indicator for the exploration of novel therapeutic strategies for cognitive deficits in schizophrenia.

To develop potential therapeutic strategy for cognitive deficits in schizophrenia, it is quite important to verify the neural substrates across a range of scales from molecular and cellular to behavioral levels in preclinical models relevant to schizophrenia (42). Growing evidence indicates that excitatory and inhibitory network in the mPFC plays a pivotal role in the regulation of working memory in normal mice (22, 43-45). Decreased dendritic spine density in the dorsolateral prefrontal cortex (DLPFC) of patients with schizophrenia has been replicated in numerous studies (46-48) and proposed to play a key pathophysiological role for schizophrenia (49). The present data of dendritic spine analysis clearly show that chronic PCP administration leads to dendritic spine loss in layer 2–3 of the PL, which is consistent with previous study (32). We also found reduction in VGLUT1+ puncta in this region, suggesting that the density of excitatory inputs in this region were reduced. Previous studies show that the size of dendritic spine head is tightly coupled to synaptic function such as the size of postsynaptic density, the number of AMPA receptors, and synaptic strength (27-29). These anatomical changes found in the present study suggest that our PCP model has attenuated excitatory inputs from other cortex in the PL which mimics morphological features in patients with schizophrenia in terms of the decreases in dendritic spines (46-48, 50) and VGLUT1 (51, 52) in layer 3 of the DLPFC.

Prefrontal interneuron impairments have been hypothesized to contribute to cognitive deficits in schizophrenia (38, 53). However, causal relationship between their abnormalities and cognitive deficits were unclear. Previous study reported that repeated blockade of NMDA receptors during early adult stage reduced the density of PV neurons in the mPFC by ascribing to a downregulation of the expression levels of PV, but not to neuronal death (54, 55). In the present study, we found not only a decrease in PV density but also a decrease in PV+ puncta onto the pyramidal neuron soma, presumed PV+ basket cell terminals, in layer 2–3 of the PL. These results suggest that chronic PCP administration during early adult stage leads to hypofunction of prelimbic PV neurons in a layer specific manner, which are in line with post-mortem brain studies of schizophrenia (38, 39). It is known that disinhibition of pyramidal cells occurs due to suppression of inhibitory neurons, which may be caused by PV neuron defects (56). Also, we previously observed PCP model showed c-Fos overexpression in layer 2–3 of the PL after working memory load (18). Based on these results, we hypothesized that attenuated inhibition from PV neurons to excitatory pyramidal neurons and these structural changes may provide a potential neurobiological basis for cognitive deficits. Importantly, in addition to working memory deficit, these abnormalities in excitatory and inhibitory synapses of the PL in our PCP model were antipsychotics-resistant.

Our histological data raise the possibility that PV neuron specific perturbation in the PL may ameliorate working memory deficit in chronic PCP-treated mice. Indeed, previous studies using pharmacological (19) or genetic (57) animal models of schizophrenia have found evidence of prefrontal PV neuron defects at cellular levels, though causal relationship between their abnormalities and cognitive deficits remain to be unaddressed. In the present study, we found that prelimbic PV activation which were achieved using a Cre-dependent excitatory hM3D(Gq)-DREADD system and DCZ ameliorate working memory deficit in our PCP model. These results suggest that specific increase in prelimbic PV neuron activity is sufficient to enhance working memory in this model. In addition, chemogenetic PV activation in this region did not disturb the inhibitory effects of continuous olanzapine treatment on acute PCP-induced hyperactivity, which may be due to sustained 60–80% occupancy of striatal dopamine D2 receptor blockade. Taken together, our findings suggest that prefrontal PV activation may be a promising therapeutic strategy to improve cognitive deficits in schizophrenia when added on current antipsychotic medication.

Present study, aiming to dissect neural substrates of working memory deficit in PCP model and to develop therapeutic strategies for them, mainly focused on PV neurons in the mPFC, especially PL. PV+ basket cells targeting the perisomatic pyramidal cells exert a powerful control over the excitability of them (58, 59). PL PV+ basket cells differentiate their firing during working memory task (60). PV+ basket cells do not fire homogeneously, but individual cells are recruited or inhibited while rats performed working memory task, implying discrete contributions to different task episodes by subsets of PV+ basket cells in the PL. Although the chemogenetic approach we used in the present study may have activated individual PV neurons by decrease in their firing threshold in the PL, the underlying circuit mechanism remain unclear. In addition, PV neurons are essential for the generation of gamma oscillations (61). Neural oscillations such as gamma band in the mPFC and hippocampus have been linked to spatial working memory in rodents (62). Gamma oscillations are observed in sample (encoding) (63) and choice phase (retrieval) (64) of working memory task. Patients with schizophrenia have lower amplitudes in gamma band than healthy controls during task, and both baseline and working memory-induced gamma show strong dependence on baseline GABA level in the DLPFC (65). However, temporal involvement of neural oscillations in working memory or its impairment in PCP model remains unclear. Further understanding of the functional dynamics of PV neurons, including neuronal ensemble or oscillatory activity such as gamma band, in working memory deficit and its ameliorating effects is needed.

In conclusion, our study strengthens the understanding of the causal relationship between prefrontal PV neurons and working memory deficit in schizophrenia. Based on this hypothesis, we propose that perturbation to prefrontal PV neurons may lead to an add-on therapy to antipsychotics, which provide new insights into drug discovery targeting amelioration of cognitive deficits in schizophrenia.

## Acknowledgements

We thank Dr. Nobuyuki Kai for valuable advice for data analyses. We are also grateful to Prof. Kazuhiro Goto of Sagami Women’s University, Sagamihara, Kanagawa, Japan for helpful input regarding the behavioral paradigm. This work was supported by JSPS KAKENHI (Grant Numbers 26860942, 17K16397 to Y.A. and 19H04899 to N.O., the Smoking Research Foundation, Dokkyo International Medical Education and Research Foundation to Y.A., the Brain Science Foundation, the Naito Foundation, and the Takeda Science Foundation to N.O.

## Author contributions

Y.A designed the study. Y.A, K.A. and N.O. supervised the project. Y.A. performed all behavioral experiments and analyzed the data. C.K. and N.O. prepared viral vectors. Y.A. and Y.S. performed animal surgery. Y.A. and M.I. performed the histological experiments.

Y.A. and N.O. performed in vivo imaging. C.K. and N.O. analyzed the data of calcium imaging. Y.A., Y.U., and K.A. developed LC-MS/MS-based in vivo dopamine D2 receptor occupancy analysis. K.F. and K.T. generated and maintained the transgenic mice.

Y.A. and N.O. wrote the manuscript, with comments from all authors.

## Supplemental Methods

### Animals

For dendritic spine and synapses analysis, male mice harboring a green fluorescent protein (GFP) transgene under control of the Thy1 promoter, were specified as Thy1-GFP line O (Strain number: 007919), and were obtained from Jackson Laboratory. Hemizygous male Thy1-GFP line O mice were crossed against inbred female C57BL/6J mice from Japan Clea Co., to give birth to hemizygotes used for breeding and morphological experiments. For in vivo dopamine D2 receptor occupancy analysis, male C57BL/6J mice (8–9 weeks old) were used. Male PV-Cre mice (Strain number: 017320) were used for in vivo Ca^2+^ imaging and DREADD experiments. Mice were housed 2–4 per cage in a temperature-controlled (24 ± 1°C) and light-controlled room (light on from 7 a.m. to 7 p.m.) in plastic cages with *ad libitum* access to food and water, except for food restriction experiments in vivo Ca^2+^ imaging. PCR-based genotyping for the transgene was conducted with primers and genomic DNA extracted from biopsy specimens of mouse tail using Hot Sodium Hydroxide and Tris (HotSHOT) method to extract genomic DNA (1).

### Drug

Phencyclidine hydrochloride was dissolved in saline. The mice were treated with saline or PCP 10 mg/kg each day for 14 consecutive days. Haloperidol and olanzapine were dissolved in a 2% glacial acetic acid in sterilized water (pH adjusted to ∼5 with 1N NaOH). For tail vein administration, S(-)-raclopride (+)-tartrate salt, used as an occupancy tracer, was dissolved in saline. For LC-MS/MS analysis, S(-)-raclopride (+)- tartrate salt and (±)-metoprolol (+)-tartrate salt was dissolved in 0.1% formic acid (LC-MS grade) in water. Deschloroclozapine (DCZ), is a highly potent and selective DREADD agonist (2), was dissolved in 1–2% dimethyl sulfoxide (DMSO) in saline.

### Viral vector preparation

For expressing hM3Dq-mCherry, mCherry, or G-CaMP7 (3) in PV+ neurons, AAV9-hSyn-DIO-hM3D(Gq)-mCherry, AAV9-hSynI-DIO-mCherry, or AAV9-hSynI-DIO-G-CaMP7 was produced as described previously (4, 5) with pAAV-hSyn-DIO-hM3D(Gq)-mCherry (Addgene #44361), pAAV-hSynI-DIO-mCherry, or pAAV-hSynI-DIO-G-CaMP7 plasmid DNA, respectively. The pAAV-hSynI-DIO-mCherry or the pAAV-hSynI-DIO-G-CaMP7 was constructed by replacing the hM3D(Gq)-mCherry sequence with the mCherry or the G-CaMP7 sequence in the pAAV-hSyn-DIO-hM3D(Gq)-mCherry plasmid using the NheI and AscI sites that were blunted with the Klenow fragment of DNA polymerase I. Viral titer was determined by quantitative polymerase chain reaction.

### Surgery and drug treatment

For continuous antipsychotic treatments, osmotic minipump (Alzet, model 1002 for vehicle and haloperidol, or 2002 and 2004 for olanzapine), containing either vehicle (VEH; 2% glacial acetic acid in sterilized water), haloperidol 0.25, 0.5, and 0.75 mg/kg/day or olanzapine 3.0, 7.5, and 10 mg/kg/day, were implanted subcutaneously to provide a steady delivery rate with its flow moderator away from the incision site under avertin or sevoflurane anesthesia. Mice were placed on a heating pad for small animals to maintain their body temperature during surgery. For in vivo Ca^2+^ imaging and DREADD approaches, surgeries were performed using stereotaxic instruments (SR-5M-HT and SM-15; Narishige). Mice were anesthetized with mixture of medetomidine, midazolam, and butorphanol.

### Immunohistochemical analysis of three-dimensional, detailed, and cortical layer specific dendritic spine analysis in the prelimbic cortex

After 1-week-withdrawal from daily PCP 10 mg/kg or saline administration for 14 days, Thy1-GFP line O mice were anaesthetized with pentobarbital (100 mg/kg, i.p.) and xylazine (10 mg/kg, i.p.), transcardially perfused with 1% paraformaldehyde (PFA) in 0.1M phosphate buffer (PB) for 1 min, and then fixative containing 4% PFA and 0.125% glutaraldehyde in 0.1M PB. Brains were post-fixed with same fixative for 6 h at 4°C and then coronally sliced (100 µm thickness) using a slicer (NeoLinear Slicer MT, Dosaka EM, Japan). After washing with Tris-buffered saline (TBS) containing 0.1% Tween 20 (TBST), free-floating sections were incubated with TBS containing 5% normal goat serum (NGS) and 0.3% Triton X-100 for 1h at room temperature (RT), and then for two overnight at 4°C in the primary antibody diluted in blocking buffer. We used the following primary antibodies: chicken anti-GFP (1:2000, ab13970, abcam), rat anti-Ctip2 (1:500, ab18465, abcam), rabbit anti-Foxp2 (1:2000, ab16046, abcam). After washing with TBST, sections were incubated for 2 h at RT with goat Alexa 488-conjugated anti chick IgG (1:500, A11039), goat Alexa 594-conjugated anti-rat IgG (1:500, A11007) and goat Alexa 647-conjugated anti-rabbit IgG (1:500, A21245). After washing, sections were mounted onto glass slides.

### Immunohistochemical analysis of excitatory and inhibitory synaptic puncta analysis in the prelimbic cortex

After 1-week-withdrawal from daily PCP 10 mg/kg or saline administration for 14 days, Thy1-GFP line O mice were anaesthetized with sodium pentobarbital (100 mg/kg, i.p.) and xylazine (10 mg/kg, i.p.), and transcardially perfused with 4% PFA and 15% saturation picric acid in 0.1 M PB. For synapse staining, the whole process of this perfusion fixation procedure was quickly performed to prevent the structural changes at synapses (6). Brains were post-fixed with same fixative for 2 h at 4°C and transferred to 15% sucrose in 0.1 M PB and then immersed in 30% sucrose in 0.1 M PB for cryoprotection. 20 µm-thick sections were cut with cryostat and stored at −30°C in a solution containing 30% (v/v) ethylene glycol, 30% (v/v) glycerol and 0.1 M sodium phosphate buffer, until use. Free-floating sections were rinsed with TBST, incubated with TBS containing 5% NGS and 0.3% Triton X-100 for 1h at RT and then incubated two (VGLUT2, GAD65, CaMK2a, and PV) or three (VGLUT1) overnight at 4°C with the primary antibodies for synapse staining and rat anti-ctip2 antibody (1:500, ab18465, abcam). We used the following antibodies for synapse puncta analysis: mouse anti-VGLUT1 (1:1000, 135-511, Synaptic Systems), guinea pig anti-VGLUT2 (1:2000, AB2251, Millipore), mouse anti-GAD65 (1:1000, MAB351, Millipore), rat anti-Ctip2 (1:500, ab18465, abcam), mouse anti-CaMK2a (1:2000, 05-532), and mouse anti-parvalbumin (1:2000, P3088, Sigma-Aldrich). After washing with TBST, sections were incubated for 2 h at RT with the secondary antibodies. We used the following secondary antibodies: goat Dylight 405-conjugated anti rat IgG (1:500, 612-146-120, Rockland), goat Alexa 488-conjugated anti-mouse IgG (1:500, A11029), goat Alexa 594-conjugated anti-mouse IgG (1:500, A11032) and goat Alexa 647-conjugated anti-guinea pig IgG (1:500, A21450). After washing, sections were mounted onto glass slides.

### Image acquisition and data analysis for dendritic spines and synapses

Acquisition of fluorescence images and image analyses were conducted by an experimenter blind to test conditions. Dendritic spine analysis was performed according to previously described methods (7) with minor modifications. Confocal images were acquired with the LSM780 (Carl Zeiss) with sequential acquisition method using 20× 0.8 NA objective at 256 × 256 pixels resolution and 63× 1.4 NA objective at 1024 × 1024 pixels resolution as z-stacks and 2.5× digital zoom. Confocal images were deconvolved with the 3D blind deconvolution software AutoQuant X3 (Media Cybernetics, MD, USA) and spine analysis was performed using the semi-automated software NeuronStudio (http://research.mssm.edu/cnic/tools-ns.html). NeuronStudio classifies spines into three types (thin, mushroom, and stubby) and analyzes dendritic length, spine number, spine head diameter and spine length. For the puncta analysis of VGLUT1, VGLUT2, GAD65 and PV immunostaining, ROI of the cortical layer in the prelimbic cortex was determined by ctip2 immunoreactive layer. Confocal images were acquired with a 63× 1.4 NA objective at 1024 × 1024 pixels resolution and 3× digital zoom. Z-stack images were obtained with approximately 15 µm optical thick (e.g. 35 × 0.43 µm optical sections for VGLUT1) excluding section surface and then deconvolved with AutoQuant X3. Quantification of fluorescence intensity, puncta density and particle size in VGLUT1, VGLUT2 and GAD65 was performed using “measure”, “threshold” and “analyze particles” function of ImageJ. Quantification of them in PV was performed using “ROI manager”, “subtract background”, “measure”, “threshold” and “analyze particles” plugins.

### In vivo calcium imaging with hM3D(Gq)-DREADD

Mice were anesthetized with mixture of medetomidine, midazolam, and butorphanol and placed on a stereotaxic apparatus. The virus solution (200 nL/site) was injected into the left PL (AP: -1.8, ML: 0.3, DV: -2.2 mm from bregma) at a rate of 0.1 µL/min using a borosilicate glass capillary. Adeno-associated viruses (AAVs) AAV9-hSyn-DIO-hM3D(Gq)-mCherry (4 × 10^13^ vg/mL) and AAV9-hSyn-DIO-G-CaMP7 (4 × 10^13^ vg/mL) were infused into the target brain areas. The glass capillary was left in place for at least 10 min. After recovery, we implanted a gradient refractive index (GRIN) lens (Inscopix, diameter 0.5 mm; length: 6.1 mm) into the left PL (AP: -1.8, ML: 0.3, DV: - 1.9 mm from bregma). We confirmed spherical and chromatic aberrations of GRIN lens using 4.0 µm TetraSpeck Fluorescent Microsphere (Thermo Scientific). The chamber frame was then cemented to fix the mouse head under the microscope. Calcium imaging experiments began at least 4 weeks after the surgery to allow time for optimal viral expression. During imaging, we head-fixed awake mouse on a treadmill. We performed calcium imaging with chemogenetic manipulation by hM3D(Gq)-DREADD. The fluorescence images of mCherry were used as a marker of hM3D(Gq)-expressing PV neurons. Calcium signals from G-CaMP7-expressed PV neurons were imaged for 10 min each condition (pretreatment, vehicle, and DCZ 3 µg/kg treatment) with 15 min interval. Vehicle or DCZ were administered 10 min before imaging. The fluorescence images of G-CaMP7 were acquired at a rate of 7.5 frames/s using a custom spinning disk confocal microscope CSU-X1 (Yokogawa), a sCMOS pco.edge camera (PCO), a 10× 0.3 NA objective (OLYMPUS), 488 nm laser (Lucir). The acquired images were first spatially downsampled by a factor of 4. For tracking cells across each condition (pretreatment, vehicle, and DCZ treatment), single session movies were concatenated into one total movie containing all the imaging sessions, followed by smoothing with Gaussian blur filter. Regions of interest (ROIs) were manually using ImageJ Fiji software. Finally, cells were identified using an automated sorting system, HOTARU (High performance Optimizer to extract spike Timing And cell location from calcium imaging data via lineaR impUlse) (8, 9), and the HOTARU output data of Ca^2+^ signals (fluorescence intensity) represent Ca^2+^ activity in each time frame in each identified cell. To determine each cell’s pattern of activity across treatment conditions, the data were normalized to baseline levels of each cell (the average of Ca^2+^ signals during pretreatment), binned into 400 ms windows, and individual z-score was calculated for each cell with the mean and standard deviation of pretreatment. We averaged z-scored Ca^2+^ signals for each treatment condition. The Ca^2+^ events were extracted with the thresholds of 3 SD of baseline. After verification that both virus-driven expression and GRIN lens were located in the left PL, imaging data were analyzed.

### Delayed nonmatching-to-position (DNMTP) task with T-maze

Working memory was assessed using delayed nonmatching-to-position task (DNMTP) with T-maze which has been described previously (10), with slight modifications. Male C57BL/6J and PV-Cre mice were 7 weeks old at the beginning of the experiment and maintained at about 85% of free-feeding body weight by supplementary feeding with normal rodent diet (CE-2) in addition to the food pellets delivered in daily sessions. Water was freely available in the home cage. All behavioral analyses were recorded using a CCD camera under approximately 50-lux illumination at the center of the T-maze. Before all experiments, mice were acclimatized to a sound-attenuated room for 30 min. This task was conducted using a T-maze constructed to two side goal arms (30 × 10 × 15 cm). Two sliding guillotine doors were manually operated to keep the test mouse in the starting area (start box) or to block their entry into the left or right goal arm. The DNMTP task in T-maze was performed over four sessions: (1) adaptation, (2) forced-alternation training, (3) DNMTP training, (4) DNMTP test. In the adaptation procedure, mice were habituated to the T-maze for 2 days. Mice were first placed in the maze with their cagemates and allowed to freely explore all arms for 10 min and then placed in the maze alone for 5 min each once daily. A Food pellet (20 mg dustless precision pellets, Bio-Serv), used as reinforcer during tasks, was placed in a pellet cup at the end of each goal arm. In the forced alternation training, the mouse was forced to alternate between left and right arm to obtain the food reward. The mouse was placed in the start box with either the left and right goal arm open, and then allowed to obtain food pellets. If the mouse did not run after 30 sec, it was gently pushed to initiate movement as appropriate. The guillotine door of the start box was re-opened after a 5 sec delay, and the mouse was allowed to run into the goal arm which was in the opposite direction to that visited before. Subsequently, it was removed from the maze, and put into a holding cage. There was a 40 sec inter-trial interval (ITI), and then the next trial was started. There were 10 trials on each training day, with the direction of the forced run varying in a pseudorandom sequence manner. This forced-alternation training was followed by DNMTP training. The mouse allowed to enter the opened arm to eat a food pellet, and then were moved back to start box. After a 5 sec delay, the mouse was allowed to access both goal arms in the choice run. When the mouse selects the goal arm, which is the opposite direction to that visited before, it allowed to consume a food pellet. This session was also repeated at 10 times in a pseudorandom sequence manner and continued until a criterion of 80% correct responses on three consecutive days was achieved. After reaching the criteria, mice were treated with saline or PCP 10 mg/kg for 14 consecutive days. After a 4-day withdrawal, they were subjected to subsequent experiments. In the DNMTP test sessions, the mouse was given 4 trials at each delay (5, 15, and 30 sec) presented in a random order at 40 sec ITI. A total of 12 trials for each delay were run over the 3 or 6 consecutive days. Percent correct responses were calculated as a performance index for spatial working memory during training and test sessions.

After completion of the DNMTP test session, mice were perfused with 4% PFA. Coronal sections of the prelimbic cortex were collected for confirmation of viral transfection when appropriate.

### Chemogenetic activation of prefrontal PV neurons during behavioral tests

The procedure is described above in detail. AAV9-hSyn-DIO-mCherry (4 × 10^13^ vg/mL) or AAV9-hSyn-DIO-hM3D(Gq)-mCherry (4 × 10^13^ vg/mL) solution (200 nL/site) was injected into the bilateral PL (AP: -1.8, ML: ±0.3, DV: -2.2 mm from bregma). Test sessions of DNMTP task with T-maze and open field test were performed approximately 3–4 weeks after surgery for the experiment infusing AAVs. DCZ 3 µg/kg or vehicle was administered intraperitoneally 10–15 min before each behavioral test. After verification that virus-driven expression was located in the bilateral PL, behavioral data were analyzed.

### Open field test

All behavioral experiments were recorded with CCD camera and measured by using a video tracking software (ANY-maze; Stoelting Co., USA). The apparatus was illuminated at approximately 300 lux. Before experiments, mice were transferred to acclimatize to the experimental site for 30 min. Mice were given saline or PCP 10 mg/kg s.c., placed individually in the center of grey plastic cages (42 × 42 × 30 cm). Locomotor activity was measured for 120 min every four days (day 1, 4, 7, 10 and 13). To evaluate the effects of continuous antipsychotic treatment with clinically relevant dose regimen on PCP 3 mg/kg s.c. challenge, open field behavior was assessed 12 days after osmotic minipump implantation. The numbers and durations of the following behavioral parameters were recorded: distance traveled; the total time spent in the center zone (24 × 24 cm); the number of entries into the center zone, the total time spent in the four corner zones (9 × 9 cm at each corner); the number of entries into the corner zone.

Immediately after completion of the open field test, mice were perfused with 4% PFA. Coronal sections of the nucleus accumbens, which is an important region of acute PCP-induced locomotor activity and PCP sensitization, were collected for c-Fos immunohistochemistry.

### c-Fos immunohistochemistry

Mice were anaesthetized with sodium pentobarbital (100 mg/kg, i.p.) and xylazine (10 mg/kg, i.p.), and transcardially perfused with 4% PFA in 0.1 M PB after completion of the open field test. Free-floating sections of the nucleus accumbens were rinsed with TBST, incubated with TBS containing 5% NGS and 0.3% Triton X-100 for 1h at RT and then incubated with rabbit anti-c-Fos antibody (1:500) overnight at 4°C. After washing with TBST, sections were incubated with goat Alexa 594-conjugated anti-rabbit IgG (1:500) for 1 h at RT. The sections were treated with 4’,6-diamidino-2-phenylindole (DAPI) (1 µg/ml, Roche Diagnostics, 10236276001) and then washed with TBS. After washing, sections were mounted onto glass slides. Fluorescence images were acquired using a fluorescence microscope (BZ-X710, Keyence). Quantification of the number of c-Fos positive cells in the ROI was performed as described previously (10).

### LC-MS/MS-based in vivo dopamine D2 receptor occupancy analysis with continuous antipsychotic treatment

In vivo binding assays for dopamine D2 receptors in mouse striatum were performed modified from the method published previously (11). During a direct infusion experiment using triple quadrupole mass spectrometer, the mass spectra for S-raclopride, used as an occupancy tracer, revealed peaks at m/z 347.1. Following optimization of mass spectrometry conditions (Supplemental Table S2), the product ion m/z 112.1 was used for S-raclopride quantification for LC-MS/MS (Figure 3A).

Similarly, the product ion m/z 116.1 was used for metoprolol quantification. To optimize a tracer dose, S-raclopride levels were assessed in the striatum and cerebellum after intravenous administration of S-raclopride at doses 3, 10, and 30 µg/kg (Supplemental Table S3). At the 30 min time point, 3 µg/kg is the highest striatal/cerebellar concentration ratio. To optimize a tracer survival interval between tracer administration and animal sacrifice, S-raclopride levels were assessed at various intervals. The concentration ratio increased with time reaching a maximum at 30 min after S-raclopride administration (Supplemental Table S4). On day 3, 13, and 28 after osmotic minipump implantation, in vivo dopamine D2 receptor occupancy analysis was performed in continuous vehicle, haloperidol, and olanzapine groups. S-raclopride (occupancy tracer) and metoprolol (internal standard) were quantified using liquid chromatography-tandem mass spectrometry (LC-MS/MS). Individual mice from each group were placed in an acrylic animal holder (φ25 × 101 mm) and injected with S-raclopride 3 µg/kg (6 nmol/kg) in sterile saline via their tail vein. Thirty min after S-raclopride injection, mice were sacrificed by cervical dislocation. The brains were immediately removed, and striatum and cerebellum were rapidly dissected, weighed and stored in 1.5 mL polypropylene centrifuge tubes at -80℃ until analysis. The striatal tissue for total binding and the cerebellar tissue for non-specific binding were used. After brain tissue samples were thawed, four volumes (w/v) of 0.1% formic acid in water was added to the tubes. Tissue samples were homogenized using an ultrasonic homogenizer (Branson, Sonifier 150). For deproteinization in samples, 200 µL of acetonitrile (LC-MS grade), containing 0.1% formic acid, was added to 50 µL of tissue homogenate. The mixture was stirred and centrifuged for 16 min at 16,000 × g at 4°C. One volume of the supernatant was added to four volumes of 0.1% formic acid in water. Then, one volume of the solution was added to one volume of metoprolol solution, used as an internal standard for LC-MS/MS analysis, and 6.7 µL of the mixture was injected into the LC-MS/MS system. The HPLC-MS/MS analysis was performed by coupling a HPLC system (Eksigent ekspert ultraLC 100-XL; SCIEX) to a triple quadrupole mass spectrometer (QTRAP 5500; SCIEX), equipped with an electrospray ionization (ESI) probe on the Turbo V source (SCIEX). Chromatographic separation was performed on an ACQUITY UPLC BEH C18 Column (2.1 mm ID × 50 mm, 1.7 µm particles; Waters) at 40°C. A temperature of 10°C was maintained in the autosampler throughout experiments. Mobile phase A and B consisted of 0.1% formic acid in water and 0.1% formic acid in acetonitrile, respectively. Compounds were separated and elute from the column using following gradient system: 0–2 min 5% B, 2–7 min 5–100% B, 7–8 min 100–5% B, 8–18 min 5% B at a flow rate of 0.2 mL/min. After elution from the column, S-raclopride and metoprolol were selectively detected in positive electrospray mode using multiple reaction monitoring (MRM) mode (Supplemental Table S2). To prevent carry-over in LC-MS/MS analysis, 0.1% formic acid in water was injected between analyte injections. To calculate analyte concentration, the ion counts in the chromatograms were determined by using the quantitation procedures in Analyst software (SCIEX).

Binding potential (BP) of dopamine D2 receptors and receptor occupancy calculations were made for each mouse employing the previously used method (11-13). The ratio of the striatum minus cerebellum (an index of specific binding)/cerebellum (an index of nonspecific binding) was used to generate an index of the BP. The percent dopamine D2 receptor occupancy by continuous antipsychotic treatment was calculated as follows:

D2 receptor occupancy (%) = 100 × {(BP_vehicle_ - BP_antipsychotics_)/BP_vehicle_}

BP_antipsychotics_ stands for the average ratio in continuous antipsychotic treatment, and

BP_vehicle_ stands for the average ratio in continuous vehicle-treated mice.

### Statistical analysis

Statistical analysis was conducted using Excel (Microsoft) and SPSS Statistics software (IBM). Data were analyzed using paired t-test, Student’s t-test, and multiple-group comparisons one-way ANOVA, two-way ANOVA, or two-way repeated measures ANOVA. The Greenhouse-Geisser correction for repeated measures was applied as necessary. Bonferroni or Dunnett T3 *post hoc* test were conducted when appropriate. Cumulative frequency distribution analysis was evaluated using Kolmogorov-Smirnov test. The detailed statistics was indicated in Supplemental Table S1.

---

**Supplemental figure S1.**
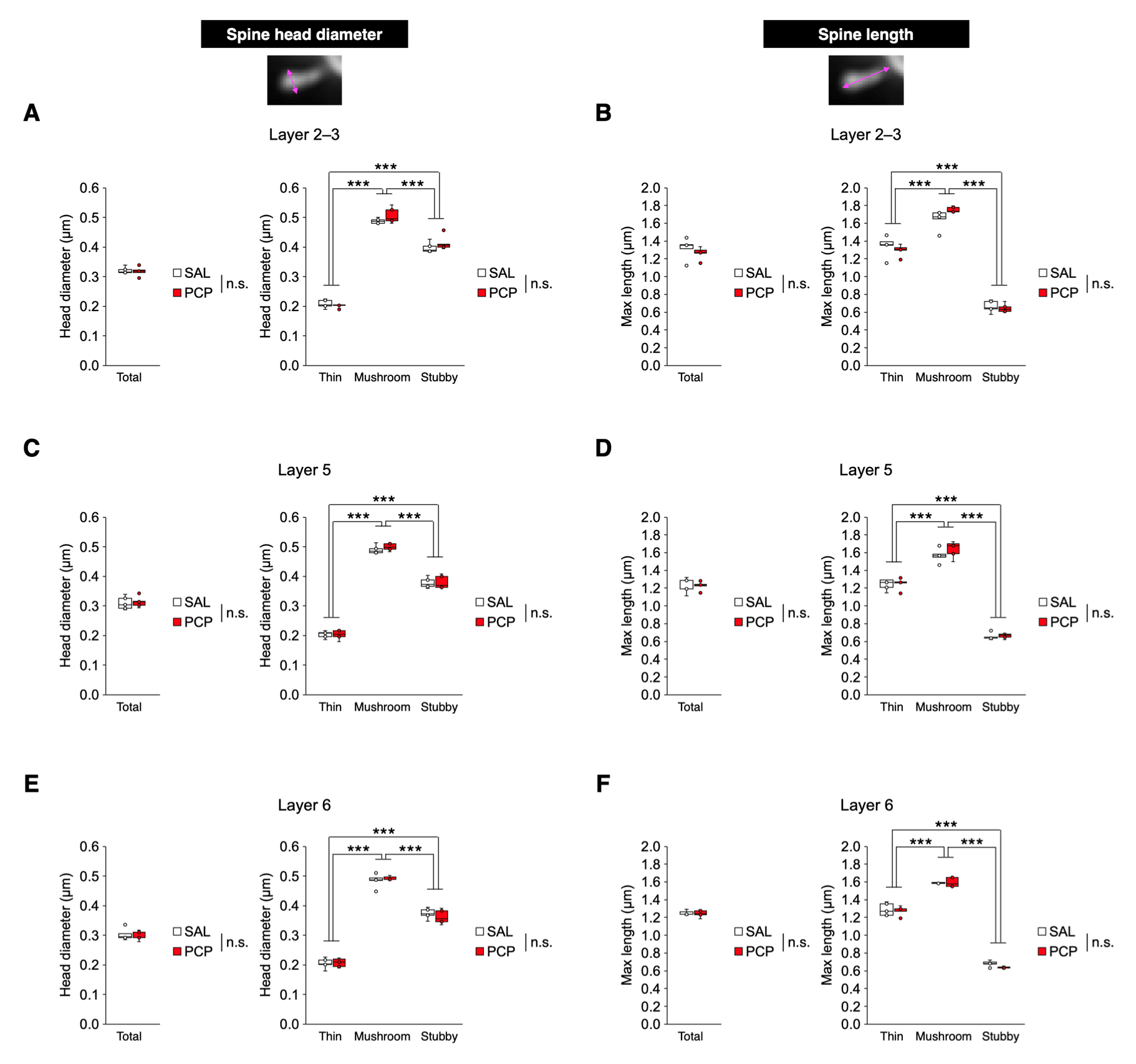
Head diameter and length of dendritic spines in the PL, related to Figure 1. **(A, C, E)** Spine head diameter in the layer 2–3 **(A**), layer 5 **(C)**, and layer 6 **(E)**. In all layers, no significant differences in head diameter of total and each spine type were observed between groups. **(B, D, F)** Spine length in the layer 2–3 **(B)**, layer 5 **(D)**, and layer 6 **(F)**. In all layers, no significant differences in spine length of total and each spine type were observed between groups. In box plots, the central mark indicates the median and the bottom and top edges of the box indicate the 25th and 75th percentiles, respectively. n.s., no significance. \*\*\**p* < 0.001.

**Supplemental figure S2.**
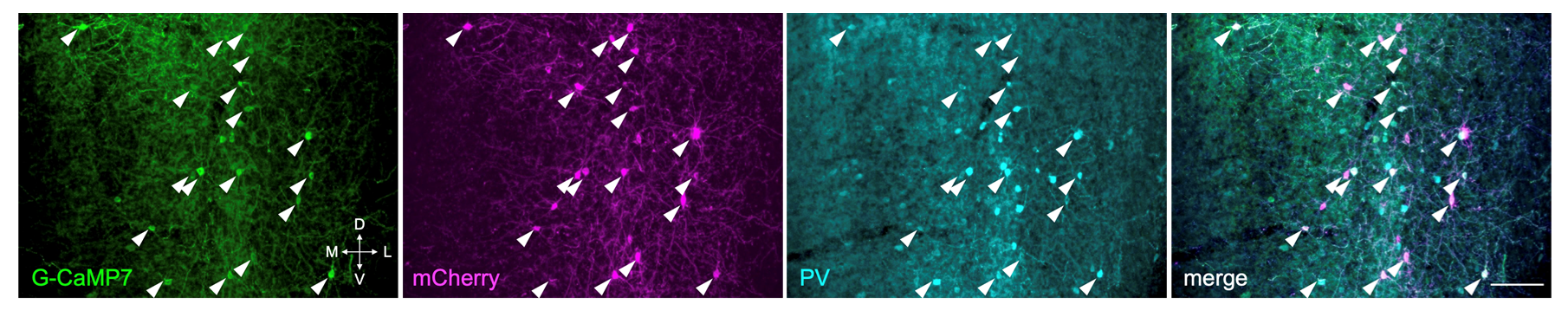
Expression of G-CaMP7 and mCherry, and PV immunostaining after calcium imaging, related to Figure 2. Representative images of G-CaMP7 (green), mCherry (magenta), and PV immunostaining (cyan) in the PL of PV-Cre mice that previously received AAV-hSyn-DIO-G-CaMP7 and AAV-hSyn-DIO-hM3D(Gq)-mCherry. White arrowheads indicate triple-labeled cells (G-CaMP7+, mCherry+, and PV+). Scale bar: 100 µm.

**Supplemental figure S3.**
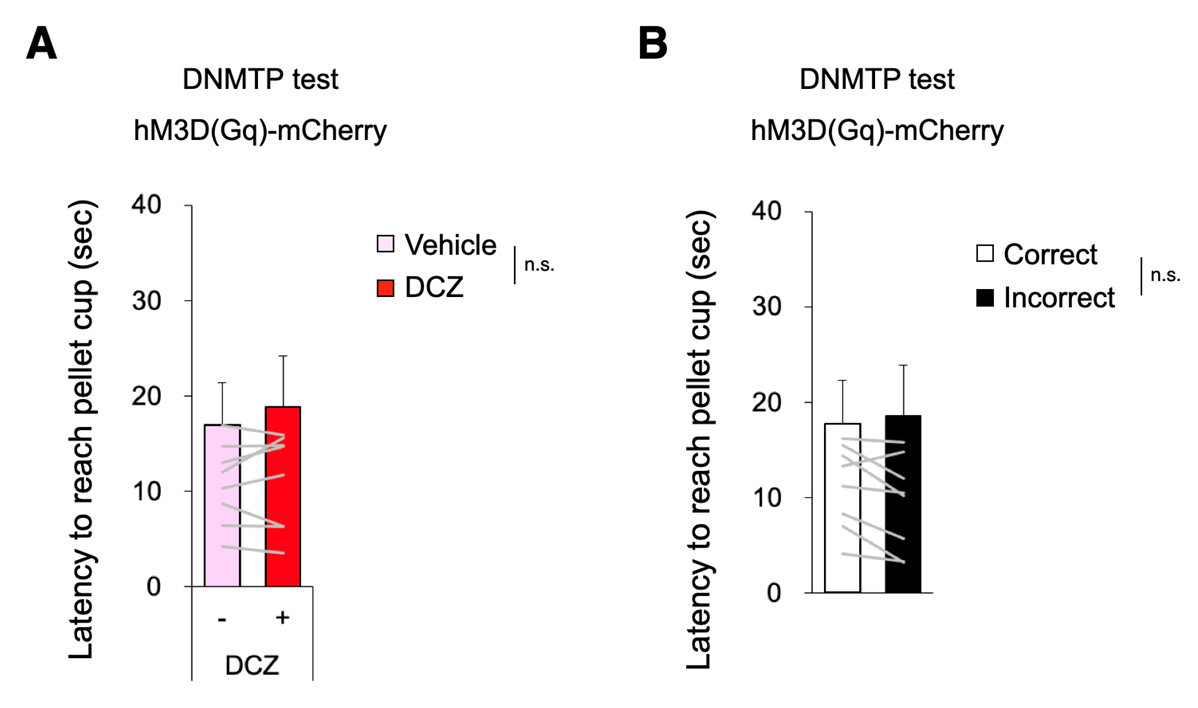
The response latency did not change with DCZ or response selections, related to Figure 2. Latency to reach pellet cup during test sessions in DNMTP task. **(A)** No significant differences in latency to reach pellet cup were observed between vehicle and DCZ treatments. **(B)** No significant differences in latency to reach pellet cup were observed between correct and incorrect responses. These results suggest response latency is not altered by treatment conditions or response selections. Data are presented as the mean ± SEM. Each line represents each mouse. n.s., no significance.

**Supplemental figure S4.**
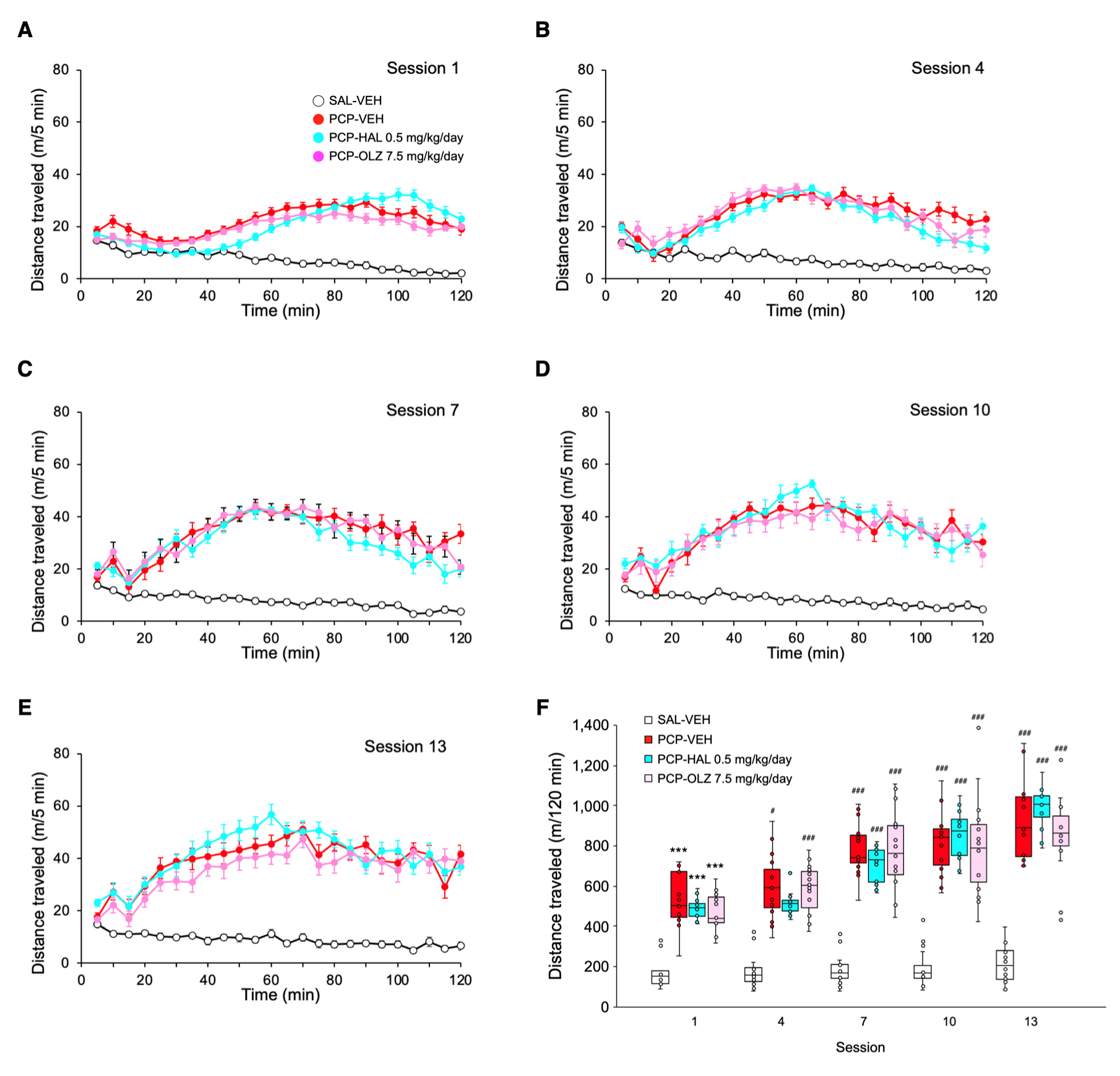
Development of PCP sensitization, related to Figure 4. **(A– E)** Sensitization of PCP-induced locomotor activity in mice. Mean distance traveled every 5 min. **(F)** Total ambulatory activity for 120 min in the open field. In box plots, the central mark indicates the median and the bottom and top edges of the box indicate the 25th and 75th percentiles, respectively. \*\*\**p* < 0.001 vs SAL-VEH in session 1. ^#^*p* < 0.05, ^###^*p* < 0.001 vs session 1 in the same group.

**Supplemental figure S5.**
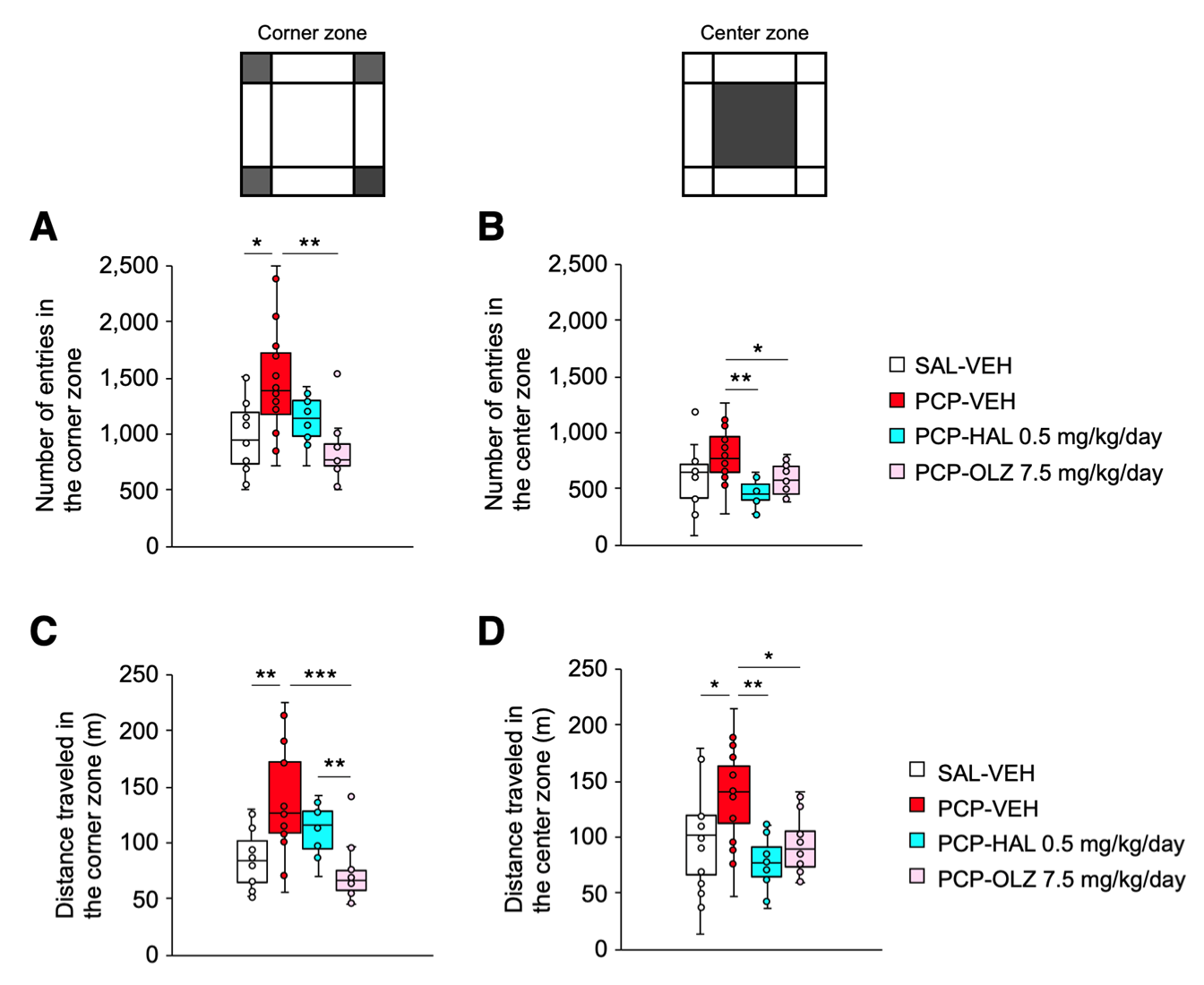
Effects of acute PCP challenge on locomotor activity pattern during continuous antipsychotic treatment, related to Figure 4. Behavioral parameters in the open field test. Number of entries into the corner **(A)** and center zone **(B)**. Total distance traveled in the corner **(C)** and center zone **(D)**. Olanzapine 7.5 mg/kg/day reversed altered behavioral patterns in chronic PCP-treated mice during open field test. In box plots, the central mark indicates the median and the bottom and top edges of the box indicate the 25th and 75th percentiles, respectively. \**p* < 0.05, \*\**p* < 0.01, and \*\*\**p* < 0.001.

**Supplemental figure S6.**
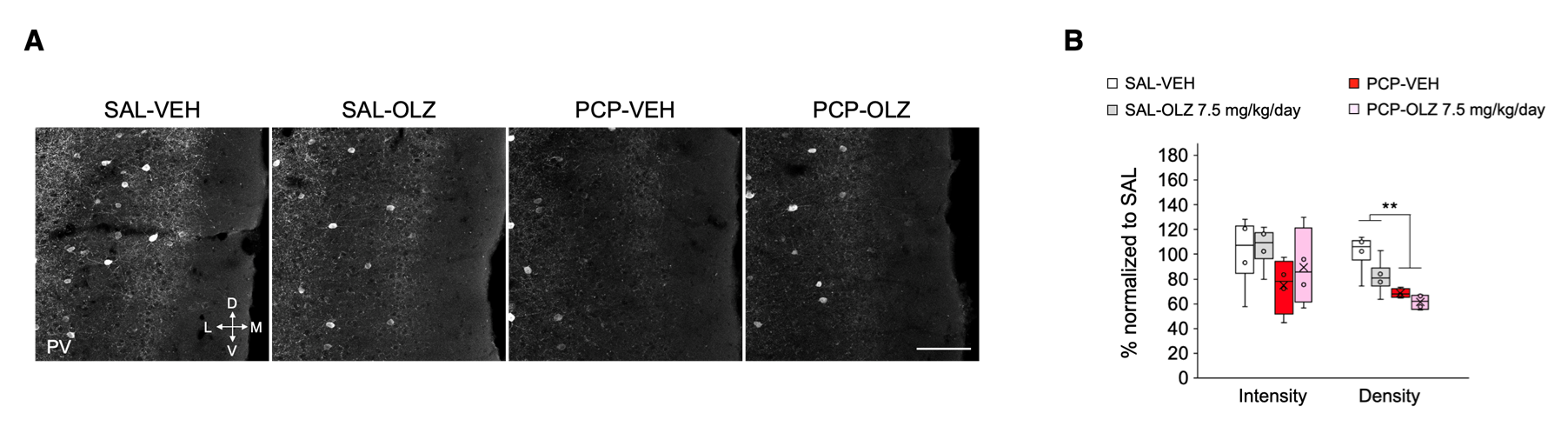
Effects of chronic PCP administration and continuous olanzapine treatment on PV neuron density in the PL, related to Figure 4. **(A)** Representative examples of immunohistochemical images. Scale bar: 100 µm. **(B)** Chronic PCP treatment significantly decreased the density of PV immunoreactive cells compared to chronic saline group. Olanzapine 7.5 mg/kg/day for 2 weeks did not affect PV neuron density both in chronic saline and PCP groups. Density: quantitative analysis of PV immunoreactive cell density. Intensity: quantitative analysis of fluorescent intensity of PV immunoreactivity. In box plots, the central mark indicates the median and the bottom and top edges of the box indicate the 25th and 75th percentiles, respectively. \*\**p* < 0.01.

**Supplemental figure S7.**
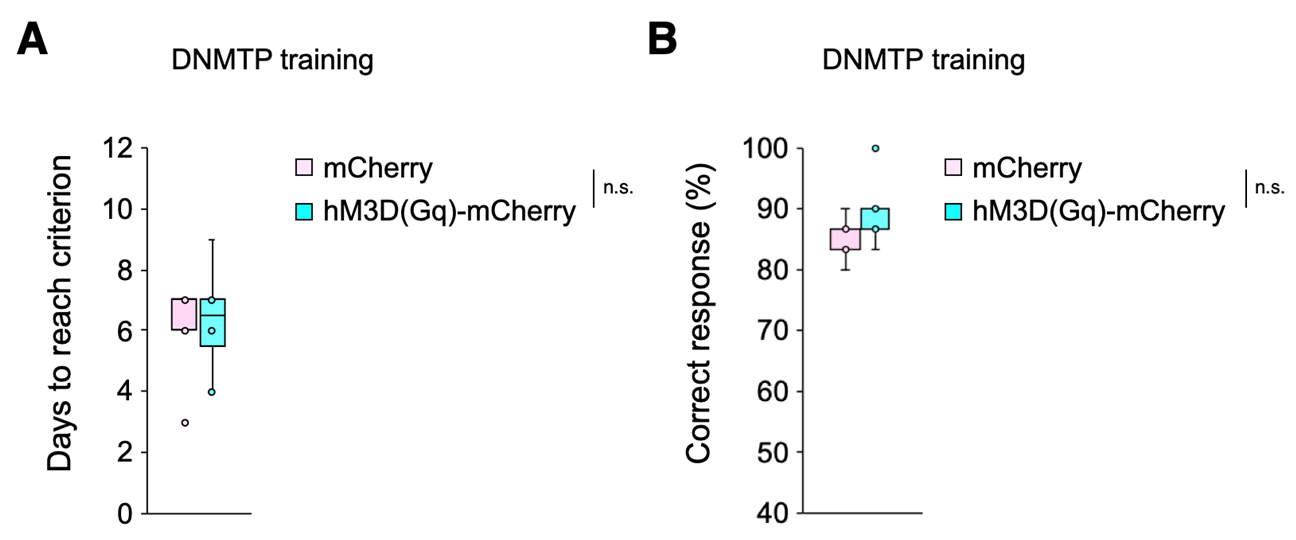
Days to reach criterion and correct responses during DNMTP training phase, related to Figure 5. Before repeated administration of PCP, no significant differences in days to reach criterion **(A)** and correct responses **(B)** were observed between mCherry- and hM3D(Gq)-mCherry-expressing mice. In box plots, the central mark indicates the median and the bottom and top edges of the box indicate the 25th and 75th percentiles, respectively. n.s., no significance.

**Supplemental figure S8.**
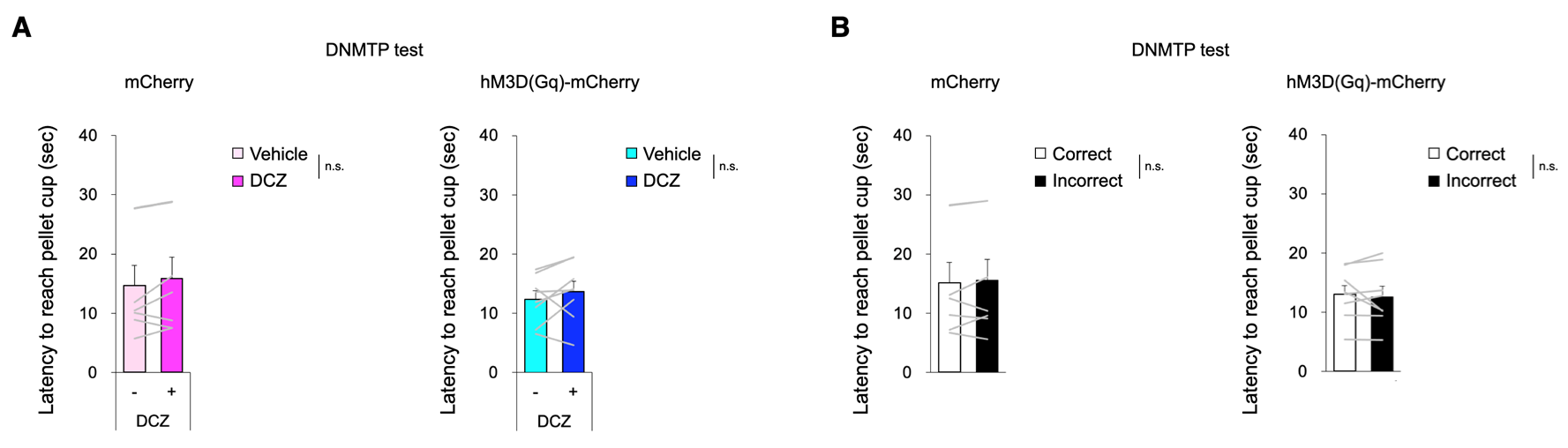
The response latency did not change with DCZ or response selections, related to Figure 5. Latency to reach pellet cup during test sessions in DNMTP task. **(A)** Both in mCherry- and hM3D(Gq)-expressing mice, no significant differences in latency to reach pellet cup were observed between vehicle and DCZ treatments. **(B)** Both in mCherry- and hM3D(Gq)-expressing mice, no significant differences in latency to reach pellet cup were observed between correct and incorrect responses. These results suggest response latency is not altered by treatment conditions or response selections. Data are presented as the mean ± SEM. Each line represents each mouse. n.s., no significance.

**Supplemental figure S9.**
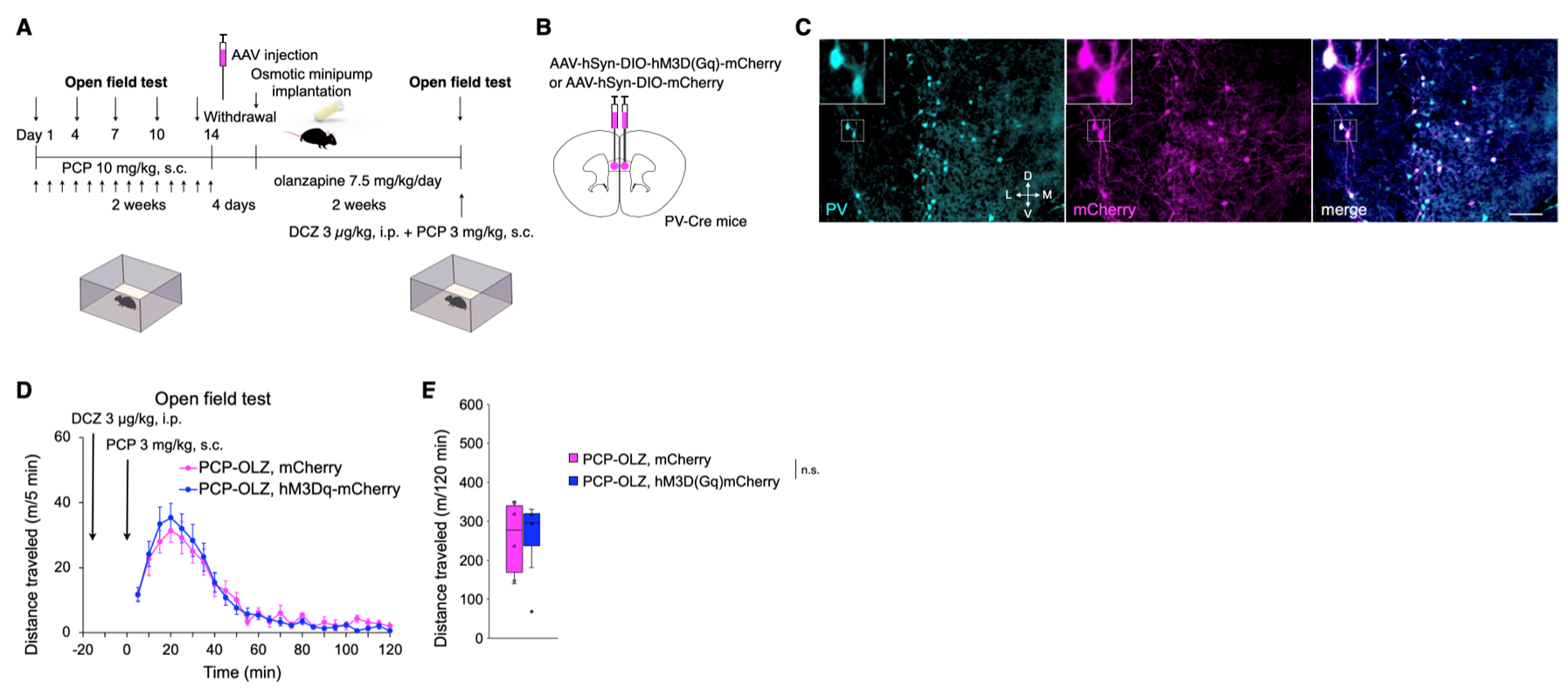
Effects of chemogenetic prefrontal PV activation on acute PCP challenge with continuous olanzapine treatment, related to Figure 5. **(A)** Schematic timelines. **(B)** Schema of bilateral viral delivery of AAV-hSyn-DIO-mCherry or AAV-hSyn-DIO-hM3D(Gq)-mCherry to PL. **(C)** Representative images of PV immunostaining (cyan) and viral expression (magenta) in the PL. Scale bar: 100 µm. **(D)** Mean distance traveled every 5 min for both mCherry- and hM3D(Gq)-expressing mice given continuous olanzapine treatment. **(E)** Total distance traveled in 120 min. In box plots, the central mark indicates the median and the bottom and top edges of the box indicate the 25th and 75th percentiles, respectively. n.s., no significance.

**Supplemental Table S1.** Statistical analysis related to Figures and Supplemental Figures.

|  |  |
| --- | --- |
| Figure 1C | Left: Student's *t*-test, Saline: $n = 5$, PCP: $n = 5$, $t_{(8)} = 3.534$, $p < 0.01$.<br>Middle: Two-way ANOVA followed by Bonferroni post hoc test, Model: $F(1, 24) = 21.255$, $p < 0.001$, Spine type: $F(2, 24) = 50.279$, $p < 0.001$, Model x Spine type interaction: $F(2, 24) = 2.527$, $p = 0.101$, Thin vs. Mushroom: $p < 0.001$, Thin vs. Stubby: $p < 0.001$.<br>Right: Kolmogorov-Smirnov $D = 0.034$, $p = 0.138$. |
| Figure 1D | Left: Student's *t*-test, Saline: $n = 5$, PCP: $n = 5$, $t_{(8)} = 1.993$, $p = 0.081$.<br>Middle: Two-way ANOVA followed by Bonferroni post hoc test, Model: $F(1, 24) = 4.213$, $p = 0.051$, Spine type: $F(2, 24) = 127.412$, $p < 0.001$, Model x Spine type interaction: $F(2, 24) = 0.590$, $p = 0.562$, Thin vs. Mushroom: $p < 0.001$, Thin vs. Stubby: $p < 0.001$.<br>Right: Kolmogorov-Smirnov $D = 0.017$, $p = 0.663$. |
| Figure 1E | Left: Student's *t*-test, Saline: $n = 5$, PCP: $n = 5$, $t_{(8)} = 0.206$, $p = 0.842$.<br>Middle: Two-way ANOVA followed by Bonferroni post hoc test, Model: $F(1, 24) = 0.048$, $p = 0.828$, Spine type: $F(2, 24) = 188.915$, $p < 0.001$, Model x Spine type interaction: $F(2, 24) = 0.233$, $p = 0.794$, Thin vs. Mushroom: $p < 0.001$, Thin vs. Stubby: $p < 0.001$.<br>Right: Kolmogorov-Smirnov $D = 0.022$, $p = 0.521$. |
| Figure 1G | Student's *t*-test, Saline: $n = 5$, PCP: $n = 5$.<br>VGLUT1: Intensity, $t_{(8)} = 1.605$, $p = 0.147$, Density, $t_{(8)} = 2.461$, $p < 0.05$, Size, $t_{(8)} = 1.557$, $p = 0.158$. VGLUT2: Intensity, $t_{(8)} = 1.112$, $p = 0.299$, Density, $t_{(8)} = 1.145$, $p = 0.285$, Size, $t_{(8)} = 0.510$, $p = 0.624$.<br>Bottom: Student's *t*-test. VGLUT1: Intensity, $t_{(8)} = 0.980$, $p = 0.356$, Density, $t_{(8)} = 1.280$, $p = 0.236$, Size, $t_{(8)} = 0.881$, $p = 0.404$, VGLUT2: Intensity, $t_{(8)} = 0.322$, $p = 0.756$, Density, $t_{(8)} = 0.169$, $p = 0.870$, Size, $t_{(8)} = 1.242$, $p = 0.249$. |
| Figure 1I | Student's *t*-test, Saline: $n = 5$, PCP: $n = 5$.<br>GAD65 in layer 2–3: Intensity, $t_{(8)} = 0.612$, $p = 0.558$, Density, $t_{(8)} = 0.720$, $p = 0.492$, Size, $t_{(8)} = 0.540$, $p = 0.604$. PV in layer 2–3: Intensity, $t_{(8)} = 1.770$, $p = 0.115$, Per soma, $t_{(8)} = 2.472$, $p < 0.05$, Size, $t_{(8)} = 0.349$, $p = 0.736$.<br>GAD65 in layer 5: Intensity, $t_{(8)} = 0.497$, $p = 0.633$, Density, $t_{(8)} = 0.428$, $p = 0.680$, Size, $t_{(8)} = 1.990$, $p = 0.082$, PV in layer 5: Intensity, $t_{(8)} = 1.013$, $p = 0.341$, Per soma, $t_{(8)} = 0.988$, $p = 0.352$, Size, $t_{(8)} = 1.381$, $p = 0.205$. |
| Figure 2E | left: One-way repeated measures ANOVA followed by Bonferroni post hoc test, 20 cells obtained from 3 mice, Treatment: $F(1.159, 22.012) = 49.924$, $p < 0.001$, DCZ vs Pretreatment: $p < 0.001$, DCZ vs VEH: $p < 0.001$.<br>right: One-way repeated measures ANOVA followed by Bonferroni post hoc test, 20 cells obtained from 3 mice, Treatment: $F(1.026, 19.498) = 36.242$, $p < 0.001$, DCZ vs Pretreatment: $p < 0.001$, DCZ vs VEH: $p < 0.001$. |
| Figure 2J | Two-way repeated measures ANOVA, $n = 9$, Delay x Drug interaction: $F(2, 14) = 0.398$, $p = 0.679$, Delay: $F(2, 14) = 0.297$, $p = 0.748$, Drug: $F(1, 7) = 16.568$, $p < 0.01$. |
| Figure 4C | One-way ANOVA followed by Dunnett T3 post hoc test, SAL-VEH: $n = 16$, PCP-VEH: $n = 15$, PCP-HAL: $n = 12$, PCP-OLZ: $n = 15$, Distance: $F(3, 57) = 8.476$, $p < 0.01$, PCP-VEH vs SAL-VEH: $p < 0.05$, PCP-HAL vs SAL-VEH: $p = 0.119$, PCP-VEH vs PCP-OLZ: $p < 0.01$ |
| Figure 4E | NAc core: One-way ANOVA followed by Dunnett T3 post hoc test, SAL-VEH: $n = 4$, PCP-VEH: $n = 4$, PCP-OLZ: $n = 4$, Distance: $F(2, 11) = 4.490$, $p = 0.044$, PCP-VEH vs SAL-VEH: $p = 0.196$, PCP-VEH vs PCP-OLZ: $p = 0.264$.<br>NAc shell: One-way ANOVA followed by Dunnett T3 post hoc test, Distance: $F(2, 11) = 22.871$, $p < 0.001$, PCP-VEH vs SAL-VEH: $p < 0.05$, PCP-VEH vs PCP-OLZ: $p < 0.05$. |
| Figure 4G | One-way ANOVA, SAL-VEH: $n = 8$, PCP-VEH: $n = 8$, PCP-OLZ: $n = 8$, Days to reach criterion: $F(2, 23) = 1.040$, $p = 0.371$. |
| Figure 4H | One-way ANOVA, SAL-VEH: $n = 8$, PCP-VEH: $n = 8$, PCP-OLZ: $n = 8$, Percent correct responses: $F(2, 23) = 1.781$, $p = 0.193$. |
| Figure 4I | Two-way repeated measures ANOVA followed by Bonferroni post hoc test, SAL-VEH: $n = 8$, PCP-VEH: $n = 8$, PCP-OLZ: $n = 8$, Delay x Group interaction: $F(2.835, 29.772) = 1.019$, $p = 0.395$, Delay: $F(1.418, 29.772) = 6.974$, $p < 0.01$, Group: $F(2, 21) = 7.363$, $p < 0.01$, PCP-VEH vs SAL-VEH: $p < 0.01$, PCP-VEH vs PCP-OLZ: $p = 1.000$, SAL-VEH vs SAL-OLZ: $p < 0.05$. |
| Figure 4L | Two-way ANOVA followed by Bonferroni post hoc test, SAL-VEH: $n = 4$, SAL-OLZ: $n = 4$, PCP-VEH: $n = 4$, PCP-OLZ: $n = 4$.<br>Intensity: Model x Treatment interaction: $F(1, 12) = 3.916$, $p = 0.071$, Model: $F(1, 12) = 2.869$, $p = 0.116$, Treatment: $F(1, 12) = 4.045$, $p = 0.067$.<br>Density: Model x Treatment interaction: $F(1, 12) = 0.577$, $p = 0.462$, Model: $F(1, 12) = 4.807$, $p < 0.05$, Treatment: $F(1, 12) = 1.877$, $p = 0.196$.<br>Size: Model x Treatment interaction: $F(1, 12) = 3.916$, $p = 0.071$, Model: $F(1, 12) = 0.035$, $p = 0.855$, Treatment: $F(1, 12) = 0.017$, $p = 0.898$. |
| Figure 4M | Two-way ANOVA followed by Bonferroni post hoc test, SAL-VEH: $n = 4$, SAL-OLZ: $n = 4$, PCP-VEH: $n = 4$, PCP-OLZ: $n = 4$.<br>Intensity: Model x Treatment interaction: $F(1, 12) = 1.017$, $p = 0.333$, Model: $F(1, 12) = 8.131$, $p < 0.05$, Treatment: $F(1, 12) = 1.022$, $p = 0.332$.<br>Per soma: Model x Treatment interaction: $F(1, 12) = 0.021$, $p = 0.887$, Model: $F(1, 12) = 26.950$, $p < 0.001$, Treatment: $F(1, 12) = 4.067$, $p = 0.067$.<br>Size: Model x Treatment interaction: $F(1, 12) = 0.225$, $p = 0.643$, Model: $F(1, 12) = 16.261$, $p < 0.01$, Treatment: $F(1, 12) = 0.028$, $p = 0.871$. |
| Figure 5D | mCherry: Two-way repeated measures ANOVA, $n = 7$, Delay x Drug interaction: $F(2, 12) = 0.475$, $p = 0.633$, Delay: $F(2, 12) = 1.779$, $p = 0.210$, Drug: $F(1, 6) =$ |
|  | 0.077, $p = 0.791$.<br>hM3D(Gq)-mCherry: Two-way repeated measures ANOVA, $n = 8$, Delay x Drug interaction: $F(2, 14) = 1.790$, $p = 0.203$, Delay: $F(2, 14) = 1.208$, $p = 0.328$, Drug: $F(1, 7) = 44.569$, $p < 0.001$. |
| Supplemental Figure S1A | Left: Student's $t$-test, Saline: $n = 5$, PCP: $n = 5$, $t_{(8)} = 0.351$, $p = 0.735$.<br>Right: Two-way ANOVA followed by Bonferroni post hoc test, Model: $F(1, 24) = 1.939$, $p = 0.177$, Spine type: $F(2, 24) = 683.451$, $p < 0.001$, Model x Spine type interaction: $F(2, 24) = 1.395$, $p = 0.267$, Thin vs. Mushroom: $p < 0.001$, Thin vs. Stubby: $p < 0.001$, Mushroom vs. Stubby: $p < 0.001$. |
| Supplemental Figure S1B | Left: Student's $t$-test. $t_{(8)} = 0.885$, $p = 0.402$. Saline: $n = 5$, PCP: $n = 5$<br>Right: Two-way ANOVA followed by Bonferroni post hoc test, Model: $F(1, 24) = 0.221$, $p = 0.642$, Spine type: $F(2, 24) = 452.660$, $p < 0.001$, Model x Spine type interaction: $F(2, 24) = 2.800$, $p = 0.081$, Thin vs. Mushroom: $p < 0.001$, Thin vs. Stubby: $p < 0.001$, Mushroom vs. Stubby: $p < 0.001$. |
| Supplemental Figure S1C | Left: Student's $t$-test, Saline: $n = 5$, PCP: $n = 5$, $t_{(8)} = 0.337$, $p = 0.745$.<br>Right: Two-way ANOVA followed by Bonferroni post hoc test, Model: $F(1, 24) = 0.476$, $p = 0.497$, Spine type: $F(2, 24) = 836.951$, $p < 0.001$, Model x Spine type interaction: $F(2, 24) = 0.225$, $p = 0.800$, Thin vs. Mushroom: $p < 0.001$, Thin vs. Stubby: $p < 0.001$, Mushroom vs. Stubby: $p < 0.001$. |
| Supplemental Figure S1D | Left: Student's $t$-test, Saline: $n = 5$, PCP: $n = 5$, $t_{(8)} = 0.148$, $p = 0.886$.<br>Right: Two-way ANOVA followed by Bonferroni post hoc test, Model: $F(1, 24) = 1.491$, $p = 0.234$, Spine type: $F(2, 24) = 528.342$, $p < 0.001$, Model x Spine type interaction: $F(2, 24) = 0.741$, $p = 0.487$, Thin vs. Mushroom: $p < 0.001$, Thin vs. Stubby: $p < 0.001$, Mushroom vs. Stubby: $p < 0.001$. |
| Supplemental Figure S1E | Left: Student's $t$-test, Saline: $n = 5$, PCP: $n = 5$, $t_{(8)} = 0.079$, $p = 0.939$.<br>Right: Two-way ANOVA followed by Bonferroni post hoc test, Model: $F(1, 24) = 0.048$, $p = 0.828$, Spine type: $F(2, 24) = 188.915$, $p < 0.001$, Model x Spine type interaction: $F(2, 24) = 0.233$, $p = 0.794$, Thin vs. Mushroom: $p < 0.001$, Thin vs. Stubby: $p < 0.001$, Mushroom vs. Stubby: $p < 0.001$. |
| Supplemental Figure S1F | Left: Student's $t$-test, Saline: $n = 5$, PCP: $n = 5$, $t_{(8)} = 0.244$, $p = 0.813$.<br>Right: Two-way ANOVA followed by Bonferroni post hoc test, Model: $F(1, 24) = 1.252$, $p = 0.274$, Spine type: $F(2, 24) = 1094.593$, $p < 0.001$, Model x Spine type interaction: $F(2, 24) = 0.722$, $p = 0.496$, Thin vs. Mushroom: $p < 0.001$, Thin vs. Stubby: $p < 0.001$, Mushroom vs. Stubby: $p < 0.001$. |
| Supplemental Figure S3A | Paired $t$-test, Vehicle: $n = 9$, DCZ: $n = 9$ $t_{(8)} = 0.846$, $p = 0.422$. |
| Supplemental Figure S3B | Paired $t$-test, Correct: $n = 9$, Incorrect: $n = 9$ $t_{(8)} = 2.204$, $p = 0.059$. |
| Supplemental Figure S4F | Two-way repeated measures ANOVA followed by Bonferroni post hoc test, SAL-VEH: $n = 16$, PCP-VEH: $n = 15$, PCP-HAL: $n = 12$, PCP-OLZ: $n = 15$, Group x |
|  | Session interaction: $F(9.524, 171.436) = 16.702, p < 0.001$.<br>Group in Session 1 (between subject): $F(3, 54) = 43.33, p < 0.001$, SAL-VEH vs PCP-VEH: $p < 0.001$, SAL-VEH vs PCP-HAL: $p < 0.001$, SAL-VEH vs PCP-OLZ: $p < 0.001$.<br>Session in SAL-VEH (within subject): $F(4, 216) = 0.61, p = 0.654$.<br>Session in PCP-VEH (within subject): $F(4, 216) = 58.12, p < 0.001$, Session 4 vs Session 1: $p < 0.05$, Session 7 vs Session 1: $p < 0.001$, Session 10 vs Session 1: $p < 0.001$, Session 13 vs Session 1: $p < 0.001$.<br>Session in PCP-HAL (within subject): $F(4, 216) = 76.36, p < 0.001$, Session 4 vs Session 1: $p = 0.402$, Session 7 vs Session 1: $p < 0.001$, Session 10 vs Session 1: $p < 0.001$, Session 13 vs Session 1: $p < 0.001$.<br>Session in PCP-OLZ (within subject): $F(4, 216) = 55.73, p < 0.001$, Session 4 vs Session 1: $p < 0.001$, Session 7 vs Session 1: $p < 0.001$, Session 10 vs Session 1: $p < 0.001$, Session 13 vs Session 1: $p < 0.001$. |
| Supplemental Figure S5A | Welch's ANOVA followed by Dunnett T3 post hoc test, SAL-VEH: $n = 16$, PCP-VEH: $n = 15$, PCP-HAL: $n = 12$, PCP-OLZ: $n = 15$, $F(3, 30.293) = 8.183, p < 0.001$, SAL-VEH vs PCP-VEH: $p < 0.05$, PCP-HAL vs PCP-VEH: $p = 0.114$, PCP-OLZ vs PCP-VEH: $p < 0.01$. |
| Supplemental Figure S5B | One-way ANOVA followed by Bonferroni post hoc test, SAL-VEH: $n = 16$, PCP-VEH: $n = 15$, PCP-HAL: $n = 12$, PCP-OLZ: $n = 15$, $F(3, 55) = 6.173, p < 0.01$, SAL-VEH vs PCP-VEH: $p = 0.065$, PCP-HAL vs PCP-VEH: $p < 0.01$, PCP-OLZ vs PCP-VEH: $p < 0.05$. |
| Supplemental Figure S5C | Welch's ANOVA followed by Dunnett T3 post hoc test, SAL-VEH: $n = 16$, PCP-VEH: $n = 15$, PCP-HAL: $n = 12$, PCP-OLZ: $n = 15$, $F(3, 30.016) = 10.948, p < 0.001$, SAL-VEH vs PCP-VEH: $p < 0.01$, PCP-HAL vs PCP-VEH: $p = 0.347$, PCP-OLZ vs PCP-VEH: $p < 0.001$, PCP-OLZ vs PCP-HAL: $p < 0.01$. |
| Supplemental Figure S5D | One-way ANOVA followed by Bonferroni post hoc test, SAL-VEH: $n = 16$, PCP-VEH: $n = 15$, PCP-HAL: $n = 12$, PCP-OLZ: $n = 15$, $F(3, 55) = 6.753, p < 0.01$, SAL-VEH vs PCP-VEH: $p < 0.05$, PCP-HAL vs PCP-VEH: $p < 0.01$, PCP-OLZ vs PCP-VEH: $p < 0.05$. |
| Supplemental Figure S6B | Two-way ANOVA followed by Bonferroni post hoc test, SAL-VEH: $n = 4$, SAL-OLZ: $n = 4$, PCP-VEH: $n = 4$, PCP-OLZ: $n = 4$.<br>Intensity: Model x Treatment interaction: $F(1, 12) = 0.132, p = 0.723$, Model: $F(1, 12) = 2.344, p = 0.152$, Treatment: $F(1, 12) = 0.556, p = 0.470$.<br>Density: Model x Treatment interaction: $F(1, 12) = 0.775, p = 0.396$, Model: $F(1, 12) = 17.162, p < 0.01$, Treatment: $F(1, 12) = 3.826, p = 0.074$. |
| Supplemental Figure S7A | Student *t*-test, PCP-OLZ + AAV-hSyn-DIO-mCherry: $n = 7$, PCP-OLZ + AAV-hSyn-DIO-hM3D(Gq)-mCherry: $n = 8$, $t_{(13)} = 0.130, p = 0.898$. |
| Supplemental Figure S7B | Student *t*-test, PCP-OLZ + AAV-hSyn-DIO-mCherry: $n = 7$, PCP-OLZ + AAV-hSyn-DIO-hM3D(Gq)-mCherry: $n = 8$, $t_{(13)} = 2.000, p = 0.067$. |
| Supplemental Figure S8A | AAV-hSyn-DIO-mCherry: Paired *t*-test. $t_{(6)} = 1.606$, $p = 0.159$. Vehicle: $n = 7$, DCZ: $n = 7$.<br>AAV-hSyn-DIO-hM3D(Gq)-mCherry: Paired *t*-test. $t_{(7)} = 1.083$, $p = 0.315$. Vehicle: $n = 8$, DCZ: $n = 8$. |
| Supplemental Figure S8B | AAV-hSyn-DIO-mCherry: Paired *t*-test, Vehicle: $n = 7$, DCZ: $n = 7$, $t_{(6)} = 0.626$, $p = 0.554$.<br>AAV-hSyn-DIO-hM3D(Gq)-mCherry: Paired *t*-test, Vehicle: $n = 8$, DCZ: $n = 8$, $t_{(7)} = 0.573$, $p = 0.584$. |
| Supplemental Figure S9E | Student *t*-test, PCP-OLZ + AAV-hSyn-DIO-mCherry: $n = 6$, PCP-OLZ + AAV-hSyn-DIO-hM3D(Gq)-mCherry: $n = 7$, $t_{(11)} = 0.036$, $p = 0.972$. |

**Supplemental Table S2.** MRM transitions and analytical conditions.

| Analytes | MRM transitions |  |  |  | Analytical conditions |  |  |  |  |  |  |  |
| --- | --- | --- | --- | --- | --- | --- | --- | --- | --- | --- | --- | --- |
|  | (m/z) |  |  |  |  |  |  |  |  |  |  |  |
|  | Q1 | Q3 | DP | CE | EP | CXP | CUR | IS | TEM | GS1 | GS2 | CAD |
| S-raclopride | 347.1 | 112.1 | 70 | 35 | 10 | 8 | 10 | 4500 | 700 | 40 | 70 | 9 |
| metoprolol | 268.4 | 116.1 | 81 | 31 |  |  |  |  |  |  |  |  |

\*DP declustering potential; CE collision energy; EP entrance potential; CXP cell exit potential; CUR curtain gas; IS ion source voltage; TEM temperature of ion source; GS1 source gas 1; GS2 source gas 2; CAD collision activated dissociation

**Supplemental Table S3.**
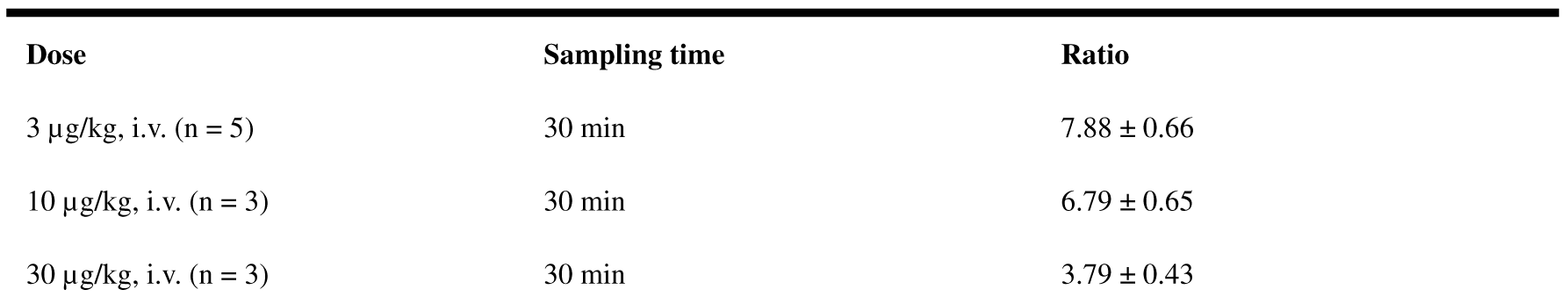
Effect of S-raclopride dose on mouse striatal/cerebellar S-raclopride concentration ratio.

| **Dose** | **Sampling time** | **Ratio** |
| --- | --- | --- |
| 3 µg/kg, i.v. (n = 5) | 30 min | 7.88 ± 0.66 |
| 10 µg/kg, i.v. (n = 3) | 30 min | 6.79 ± 0.65 |
| 30 µg/kg, i.v. (n = 3) | 30 min | 3.79 ± 0.43 |

**Supplemental Table S4.** Effect of sampling time on mouse striatal/cerebellar S-raclopride concentration ratio.

| **Dose** | **Sampling time** | **Ratio** |
| --- | --- | --- |
| 3 µg/kg, i.v. (n = 4) | 5 min | 2.69 ± 0.09 |
| 3 µg/kg, i.v. (n = 3) | 15 min | 6.98 ± 0.95 |
| 3 µg/kg, i.v. (n = 5) | 30 min | 7.88 ± 0.66 |

**Supplemental Table S5.** Dopamine D2 receptor occupancy at different doses of antipsychotics.

| **Drug** | **Doses** | **Sampling time** | **D2 receptor occupancy (%)** |
| --- | --- | --- | --- |
| Haloperidol | 0.25 mg/kg/day | Day 3 | 79.6 ± 2.95 |
|  |  | Day 13 | 48.8 ± 3.18 |
|  | 0.5 mg/kg/day | Day 3 | 80.6 ± 2.97 |
|  |  | Day 13 | 63.5 ± 3.55 |
|  | 0.75 mg/kg/day | Day 3 | 90.7 ± 1.13 |
|  |  | Day 13 | 78.3 ± 1.19 |
| Olanzapine | 3.0 mg/kg/day | Day 3 | 48.1 ± 6.49 |
|  |  | Day 13 | 48.4 ± 7.39 |
|  | 7.5 mg/kg/day | Day 3 | 80.9 ± 3.70 |
|  |  | Day 13 | 70.8 ± 2.80 |
|  |  | Day 28 | 67.2 ± 2.51 |
|  | 10 mg/kg/day | Day 3 | 79.9 ± 1.44 |
|  |  | Day 13 | 73.4 ± 2.78 |

